# Eya-controlled affinity between cell lineages drives tissue self-organization during *Drosophila* oogenesis

**DOI:** 10.1101/2022.07.28.501863

**Authors:** Vanessa Weichselberger, Patrick Dondl, Anne-Kathrin Classen

## Abstract

The cooperative morphogenesis of cell lineages underlies the development of functional units and organs. To study mechanisms driving the coordination of lineages, we investigated soma-germline interactions during oogenesis. From invertebrates to vertebrates, oocytes develop as part of a germline cyst that consists of the oocyte itself and so-called nurse cells, which feed the oocyte and are eventually removed. The enveloping somatic cells specialize to either facilitate oocyte maturation or nurse cell removal, which makes it essential to establish the right match between germline and somatic cells. We uncover that the transcriptional regulator Eya, expressed in the somatic lineage, controls bilateral cell-cell affinity between germline and somatic cells in *Drosophila* oogenesis. Employing functional studies and mathematical modelling, we show that differential affinity proportional to Eya expression and the resulting forces drive somatic cell redistribution over the germline surface and control oocyte growth to match oocyte and nurse cells with their respective somatic cells. Thus, our data demonstrate that differential affinity between cell lineages is sufficient to drive the complex assembly of inter-lineage functional units and underlies tissue self-organization during *Drosophila* oogenesis.

## Introduction

Throughout development, it is essential that different cell lineages coordinate their morphogenesis to construct functional units. This requires self-organizing mechanisms that ensure that the right cells come into contact with each other and give rise to desired shapes. Such higher-order organization emerges from simple behaviours of individual cells guided by local information ^1^. While many studies investigate self-organization within a lineage ^2, 3^, the literature is limited on functional studies elucidating mechanisms of self-organization across multiple cell lineages.

Oogenesis is a prime example of a developmental process that depends on the close interaction of two lineages. From invertebrates to vertebrates, oocytes develop within a germline cyst that is enveloped by somatic cells ^4^. The germline cyst consists of the oocyte itself and so-called nurse cells ^5–8^. The role of nurse cells is to supply the oocyte with essential materials during oogenesis, but eventually, nurse cells are removed to generate a single mature oocyte ^6, 7, 9^. The maturation of the oocyte, as well as the removal of nurse cells is strictly dependent on the cooperation with somatic cells enveloping the germline cyst. These so-called follicle cells (FCs) differentiate into diverse cell fates, which, among others, specialize to facilitate oocyte maturation by eggshell secretion or nurse cell removal by phagoptosis ^4, 7, 9–12^. Thus, it is essential that germline cells and somatic cells match each other, such that nurse cells are in contact with FCs that facilitate nurse cell removal, and the oocyte is in contact with FCs that enable oocyte maturation.

In *Drosophila* oogenesis, the establishment and maintenance of this match is a complex process that involves major cell redistributions. Oocytes develop within so-called egg chambers that consist of the germline cyst and a surrounding monolayer follicle epithelium. The germline cyst consists of 1 oocyte and 15 nurse cells, and the approximately 850 FCs differentiate into three major fates ^4, 10, 12^. Whereas the fate of germline cells is determined already prior to egg chamber assembly ^8^, FCs differentiate and specialize under the control of JAK/STAT, EGF and Notch signalling pathways during early egg chamber development ^13–20^. At the anterior tip of the egg chamber, approximately 10% of FCs differentiate into so-called anterior FCs (AFCs), which specialize to facilitate nurse cell removal and thus must cover the entire nurse cell compartment by mid-oogenesis. The remaining ∼90% of FCs differentiate into main body FCs (MBFCs) and posterior FCs (PFCs), which specialize to facilitate oocyte maturation and thus must eventually cover the entire oocyte. However, at the time point of fate specification, germline cells and FCs do not match yet: MBFCs are initially in contact with nurse cells and AFCs cover only a small proportion of the nurse cell compartment ^4, 10, 12, 21, 22^. Thus, FCs must redistribute over the germline surface to establish the right match. This redistribution must be coordinated with germline growth and changes in oocyte and nurse cell proportions, as the oocyte comprises only ∼6% of the germline (1/16^th^) when FCs are specified but makes up ∼40% of the germline in mid-oogenesis ^21^. Consequently, germline and FCs must establish the right match under constantly changing morphologies. How germline and FCs coordinate to establish inter-lineage functional units essential to produce a fertile egg is currently not understood.

## Results

### Egg chamber morphogenesis is divided into three phases with distinct soma-germline matching dynamics

To analyse the dynamics of the soma-germline matching process, we performed an in-depth quantitative description of egg chamber morphogenesis. We quantified 24 morphological parameters in egg chambers from stages 2 to 12 (Fig. 1a, Fig. S1). These parameters included among germline-, and FC-specific descriptors ^21, 23^, importantly, 10 parameters that characterized interactions between germline and FCs. To extract the dynamics of global egg chamber morphogenesis, we analysed the multidimensional dataset using UMAP (Uniform Manifold Approximation and Projection) ^24^ (Fig. 1b,c). In the UMAP projection, individual egg chambers organized along a developmental trajectory of classical egg chamber stages (Fig. 1c). Importantly, germline sizes steadily increased along the trajectory, demonstrating that germline area can be used as a continuous variable representing developmental progression (Fig. 1d). This represents an advancement over classical egg chamber staging, which is dependent on morphological features often disrupted by genetic manipulations (for example ^22^) and produces only a discrete description of a continuous process (see Supp. S1).

**Figure 1:**
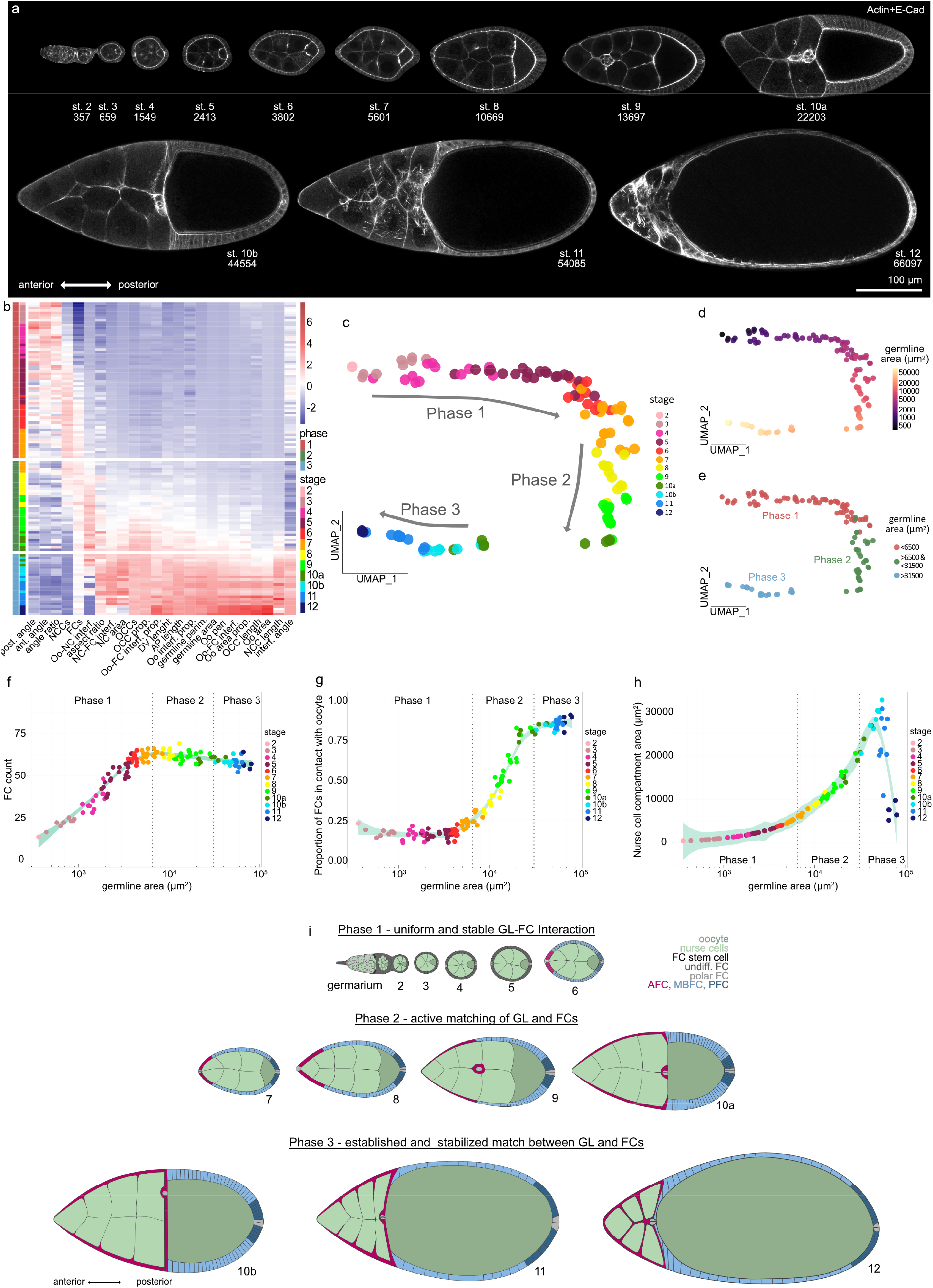
Egg chamber morphogenesis is divided into three phases with distinct soma-germline matching dynamics. **a,** Medial confocal sections of wild type (wt) egg chambers depicting developmental stages from the germarium to stage 12, stained for E-Cad and F-Actin. Numbers denote germline areas in µm^2^. **b,** Heatmap of the quantified 24 morphological parameters (see Fig. S1). Each row represents an individual egg chamber, with increasing germline sizes from top to bottom. Breaks separate morphogenetic phases. **c,** UMAP of multidimensional quantification of egg chamber morphogenesis. Egg chambers are coloured according to their respective developmental stage. Note that the developmental trajectory is subdivided into three phases. **d**, UMAP with germline areas visualized. **e**, UMAP with egg chambers assigned to the three phases based on their germline size. **f,** Follicle cell (FC) count as a function of germline area. **g,** Proportion of FCs in contact with the oocyte as a function of germline area. **h,** Nurse cell compartment area as a function of germline area. Dotted lines mark germline sizes at the transition between two phases. All curves are LOESS fitted with a 95% CI area. n=126 egg chambers. See Supp. File S2 for detailed statistical information. **i,** Illustration of the three morphogenetic phases of *Drosophila* egg chamber development.

The UMAP analysis revealed that egg chamber morphogenesis is subdivided into 3 phases. To characterize these phases, we assigned egg chambers based on their germline size to their respective phase and analysed individual morphological parameters as a function of germline size (Fig. 1e). As the differentiation of FCs into AFCs, MBFCs and PFCs at stage 6, is a crucial step for egg chamber development and coincides with an arrest of mitotic divisions ^10, 14, 15^, we analyzed the number of FCs (Fig. 1f). We found that FCs cease to multiply, and thus receive the information with which germline cell they must match, at the end of phase 1 (Fig. 1f,i). To understand the dynamics of FC redistribution that matches FCs and germline cells, we analysed the proportion of FCs that was in contact with the oocyte (Fig. 1g). Throughout phase 1, 17 ± 3% of FCs were in contact with the oocyte. By phase 3, this proportion had increased to 84 ± 3%. Thus, the redistribution of FCs and therefore the active matching between FCs and germline cells is executed during phase 2 (Fig. 1i). This suggests that the right match is essential for phase 3. Indeed, nurse cell dumping, during which nurse cell volume decreases massively, has been shown to depend on the right match and takes place during phase 3 ^25, 26^ (Fig. 1h,i).

Taken together, our multidimensional analysis reveals that global egg chamber morphogenesis is subdivided into three phases, which correlate with three distinct soma-germline interaction dynamics (Fig. 1i).

### FC distribution over germline cells co-evolve with Eya expression patterns

The matching of FCs and germline cells must be coordinated by interactions at the soma-germline interface (Fig. 2a). Cell-cell interactions are controlled by surface tension at the interface ^2, 3^ and can be described along a spectrum of cell-cell affinity to cell-cell repulsion, where affinity causes an increase in contact size between cells and repulsion a decrease (Fig. 2b). We therefore quantified the size of apical FC surfaces, which are in contact with the germline, to characterize how FCs interact with germline cells (Fig. 2c,d,f). We found that throughout phase 1, contact areas of FCs were similar, suggesting that coverage of the available germline surface was evenly distributed among all FCs. However, during phase 2, a gradient in contact areas developed, with AFCs increasing their apical contact surfaces more rapidly than the remaining FCs. The gradient resolved by phase 3 and resulted in a segregation of FCs with large contact surfaces over nurse cells and FCs with comparatively small surfaces over the oocyte.

**Figure 2:**
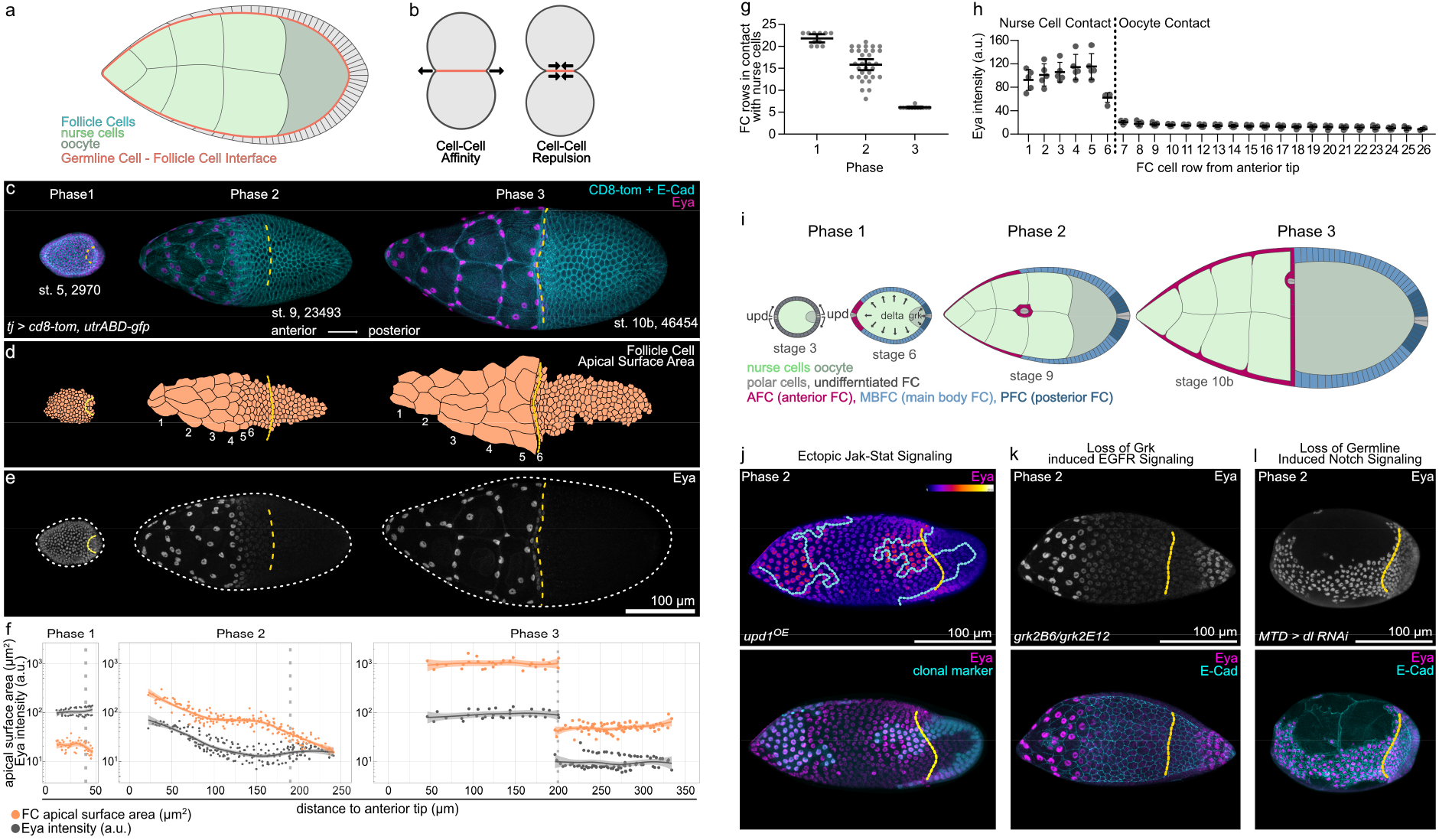
FC contact surfaces with germline cells correlate with Eya expression patterns. **a,** Illustration of the germline-soma interface. **b,** Illustration of cell-cell affinity and cell-cell repulsion. **c,** Max projections of egg chambers expressing *CD8-tom* (membrane) and *utrABD-gfp* (actin) under the control of *tj-GAL4* (FC driver), stained for E-Cadherin (E-cad) and Eyes absent (Eya). Numbers denote germline area in µm^2^. **d,** Segmented apical contact surfaces of FC with the germline of ECs shown in (c). Numbers denote FC rows from anterior to posterior. **e,** Max projections of ECs from (c), depicting expression of Eya. White dotted lines mark EC outlines. **f,** FC apical surface areas (orange) and Eya levels (grey) as a function of their distance to the anterior tip of ECs. Dotted lines mark the oocyte-nurse cell boundaries. Stage 4 EC (phase 1, n=45 FC of 2 EC); stage 9 EC (phase 2, n=192 FC of 3 EC); stage 10b EC (phase 3, n=112 FC of 3 EC). Curves are LOESS fitted with a 95% CI area. **g,** Count of FC rows in contact with nurse cells in wt EC. Mean+95% CI, n (Phase 1 = 11 EC, Phase 2 = 32 EC, Phase 3 = 12 EC). **h,** Mean Eya fluorescence intensities in FC rows in phase 3 (stage 10a & 10b). Mean±SD, n=5 EC. Dotted line separates FC rows in contact with nurse cells from FC rows in contact with oocyte. **i,** Illustration of FC patterning. During stage 3, polar cells release Upds and induce a JAK/STAT signalling gradient in nearby FCs. During stage 6, the oocyte activates *EGFR* in oocyte-contacting cells through Grk. Dl from the germline activates Notch in FCs, which allows FCs to adopt their fate. JAK/STAT + Notch = AFC, JAK/STAT + *EGFR* + Notch = PFC, Notch-only = MBFC. **j,** Max projection of a phase 2 EC with clonal expression of *upd1* and *GFP*, stained for Eya. Ectopic JAK/STAT leads to ectopic Eya gradient in MBFCs, but not PFCs. **k,** Max projection of a phase 2 *grk(2B6) /grk(2E12)* (EGF) mutant EC, stained for Eya and E-Cad. Loss of *EGFR* signalling in PFCs leads to ectopic Eya expression. **l,** Max projection of a phase 2 EC expressing *delta(dl)*-RNAi using MTD-Gal4 (germline driver), stained for Eya and E-Cad. Loss of Notch signalling causes failure of Eya downregulation in MBFCs and PFCs. See Supp. File S2 for detailed statistical information.

The gradual increase of AFC contact surfaces during phase 2 is called AFC flattening and is specific to AFC fate ^22, 27, 28^. Previous studies suggested that the increase in AFC areas could solely be a result of AFCs being stretched within the epithelium to accommodate germline growth ^12, 21, 29, 30^. To test this idea, we reduced intra-epithelial cohesion by removing the cell-cell adhesion molecules E-Cadherin and/or N-Cadherin ^3, 22^ (Fig. S2a,b). We found that this manipulation did not disrupt AFC expansion nor flattening. Secondly, we fully uncoupled individual AFCs by limiting cellular growth via ectopic expression of *hpo* ^31^. As the germline grows, the reduced cell size of affected AFCs caused them to detach from each other. Yet, these FCs maximized contacts with nurse cells by spreading out via elaborate protrusions (Fig. S2c). We therefore propose that AFCs expand apical surfaces independent of intra-epithelial cohesion by actively and autonomously increasing their contact surface with nurse cells.

In search for a regulator of FC interaction with the germline, we identified Eya (Eyes Absent). Eya is a highly conserved transcriptional co-regulator and phosphatase, and well-characterized for its role in eye specification ^32–35^. Eya is also reported to distinguish FC fate from polar and stalk cell fate during egg chamber assembly and is used as a functionally uncharacterized marker for AFC fate ^13, 21, 36–38^. We found that Eya expression patterns appeared with similar dynamics as apical FC surface sizes, with uniform expression in phase 1, a gradient in Eya levels from anterior to posterior during phase 2 and a strict segregation of Eya-positive FCs over nurse cells and Eya-negative FCs over the oocyte by phase 3 (Fig. 2e,f, Fig. S2d,e). A cell row-wise analysis revealed that this dynamic robustly led to 6 rows of FCs in contact with nurse cells by phase 3 and confirmed that exclusively these 6 rows maintained Eya expression (Fig. 2g,h).

As Eya expression patterns track with AFC fate after phase 1 (Fig. S2f), we asked if the signalling pathways determining AFC fate also control Eya dynamics. FC specification is controlled by the Jak-Stat, EGFR and Notch signalling pathways (Fig. 2i). Specifically, polar cells at each egg chamber pole secrete the ligand Upd and thereby induce a gradient of Jak-Stat signalling in surrounding FCs. In addition, the oocyte secretes the ligand Grk, which activates EGFR signalling in posterior cells. Lastly, the germline induces Notch signalling in all FCs by providing Delta, which allows FCs to adopt the fate they were primed for by Jak-Stat and EGFR signalling. Thus, FCs with Jak-Stat and Notch activation differentiate into AFCs, FCs with Jak-Stat, EGFR and Notch activation differentiate into PFCs and FCs which solely activate Notch become MBFCs ^13–20^ (Fig. 2i). We found that Eya expression was positively regulated by an ectopic Upd-induced Jak-Stat signalling gradient (Fig. 2j, Fig. S2g), negatively regulated by EGFR activation (Fig. 2k, Fig. S2g) and that the switch from uniform levels to an anterior-posterior Eya-gradient was dependent on Notch signalling inducing FC differentiation (Fig. 2l, Fig. S2g). Thus, by the end of phase 1, Eya expression becomes dependent on AFC fate specification and therefore tracks with AFCs during phase 2 and 3.

### Eya controls the size of FC contacts with germline cells

The correlation between Eya patterns and FC contact surfaces in combination with the clear segregation of Eya-positive FCs over nurse cells and Eya-negative FCs over the oocyte made us question whether Eya played a role in soma-germline matching. To test this, we manipulated Eya expression in FC clones during phase 2, when FC-germline matching takes place. First, we ectopically expressed Eya in MBFC clones, which normally lose Eya expression and transition onto the oocyte. Ectopic Eya was sufficient to cause an increase of MBFC surfaces in contact with nurse cells, which occurred via broad apical protrusions extending towards and displacing apical surfaces of neighbouring Eya-negative MBFCs (Fig. 3a-c). Next, we reduced Eya expression in AFCs, which normally express Eya and expand their contact with nurse cells and found that clonal expression of *eya-RNAi* caused a failure of contact surface increase (Fig. 3d-f). Lastly, we ectopically expressed Eya in MBFC clones, which had already transitioned onto the oocyte and found that Eya expression had no effect on the apical surface size of FCs in contact with the oocyte (Fig. 3g-i). Thus, Eya induces FCs to expand their contact surface exclusively with nurse cells, which led us to hypothesize that Eya causes FCs to experience cell-cell affinity towards nurse cells, but not towards the oocyte.

**Figure 3:**
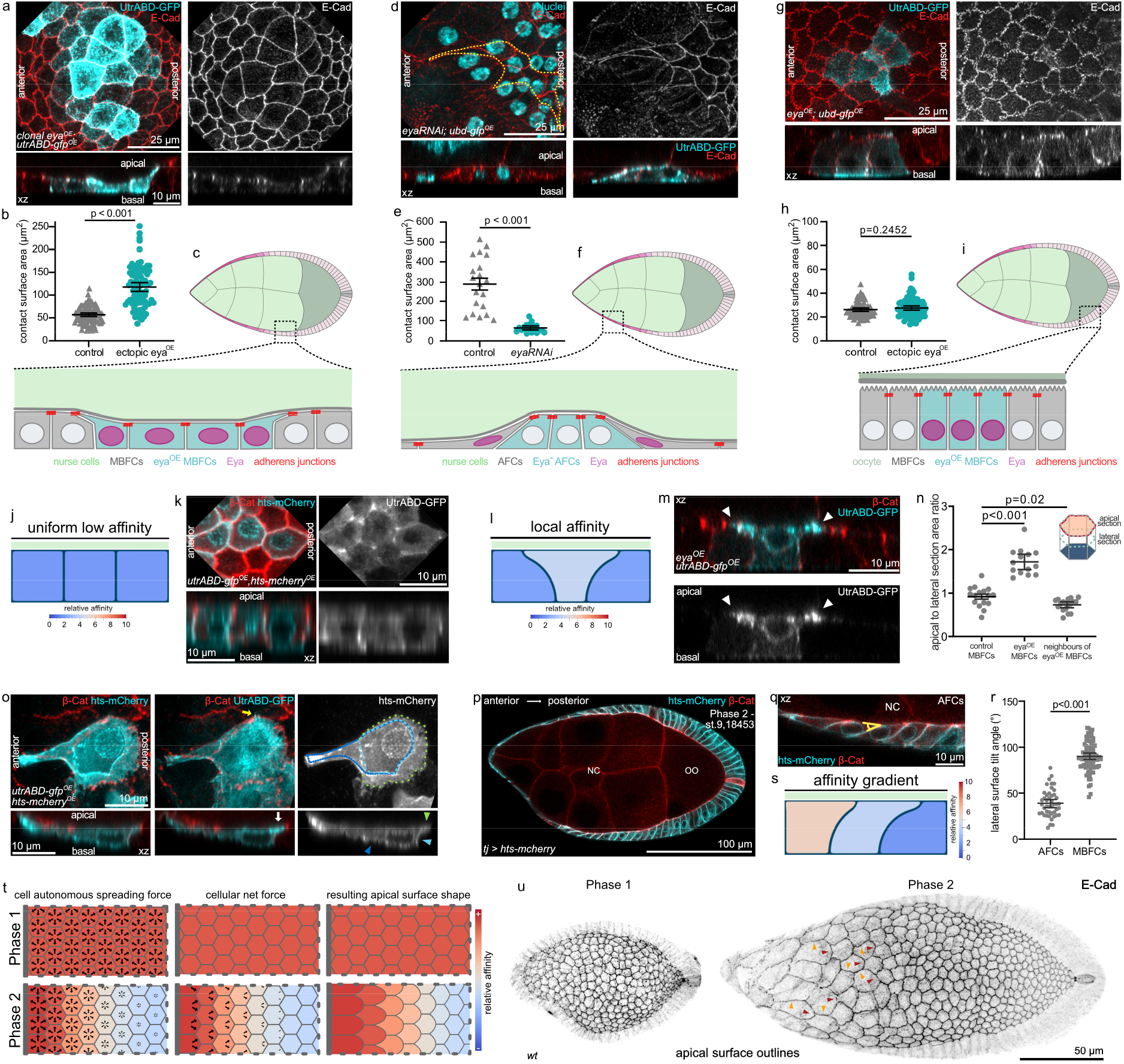
Eya expression in FCs induces Affinity for Nurse Cells. **a,** MBFCs in contact with nurse cells during phase 2 with clonal expression of *utrABD-gfp* and ectopically expressing Eya (*eya^OE^*), stained for E-Cad. Apical surface projection and xz-reslice shown. **b,** Quantification of apical contact surface areas of control MBFCs and *eya^OE^*-MBFCs in contact with nurse cells during phase 2. Mean+95% CI. Two-tailed Welch’s t-test. n (control: 71 MBFC, *eya^OE^*: 64 MBFC). **c,** Illustration of cell morphologies upon ectopic *eya^OE^* expression in MBFC clones in contact with nurse cells during phase 2. **d,** AFCs in contact with nurse cells during phase 2 with clonal expression of *eya-RNAi*, stained for E-Cad and nuclei (DAPI). Yellow line depicts clonal outline. Apical surface projection and xz-reslice shown. **e,** Quantification of apical contact surface areas of control and *eya-RNAi* AFCs during phase 2. Mean+95% CI, two-tailed Welch’s t-test, n (control: 20 AFCs, *eya-RNAi*: 20 AFCs). **f,** Illustration of cell morphologies upon *eya-RNAi* expression in AFCs during phase 2. **g,** MBFCs in contact with the oocyte during phase 2 with clonal expression of *utrABD-gfp* and *eya^OE^*, stained for E-Cad. Apical surface projection and xz-reslice shown. **h,** Quantification of apical contact surface areas of control MBFCs and MBFCs ectopically expressing *eya^OE^* in contact with the oocyte. Mean+95% CI. Unpaired Student’s t-test. n (control: 75 cells, *eya^OE^:* 83 cells). **i,** Illustration of cell morphologies upon *eya^OE^* expression in MBFC clones in contact with the oocyte during phase 2. **j,** Phase field model of 3 FCs in contact with nurse cells with low and equal affinities. **k,** MBFCs in contact with nurse cells during phase 2 with clonal expression of *utrABD-gfp* and *hts-mCherry* (membrane), stained for β-catenin (junctions). Apical surface projection and xz-reslice shown. **l,** Phase field model of 3 FCs in contact with nurse cells with the central cell developing a relatively higher affinity. **m,** MBFCs in contact with nurse cells during phase 2 with one MBFC expressing *utrABD-gfp* and *eya^OE^*, stained for β-catenin. xz-reslice shown. White arrowheads point to apical actin-rich protrusions extending towards neighbouring Eya-negative FCs. **n,** Quantification of apical to lateral area ratios (see illustration and Fig S3a) for control MBFCs, *eya^OE^*-MBFCs and direct neighbours of *eya^OE^*-MBFCs. Mean+95% CI, Welch one-way Anova with Dunnett’s T3 multiple comparisons test. n (control MBFCs: 16 cells, *eya^OE^*-MBFCs: 14 cells, neighbours: 18 cells). **o,** AFCs in phase 2 with clonal expression of *utrABD-gfp* and *hts-mcherry*, stained for β-catenin. Dotted lines outline apical (green), lateral (light blue) and basal (dark blue) surfaces. Arrowhead of same colour marks surfaces in xz. Yellow arrow points at actin-based filopodium. White arrow points at actin rich apical surface protrusion. Max projection of whole cell and xz-reslice shown. **p,** Medial confocal section of a phase 2 egg chamber expressing *hts-mCherry* under the control of *tj-GAL4* (FC driver), stained for β-cat. Germline area in µm^2^. **q,** Tilt of lateral membranes in AFCs. Angle for quantification is depicted in yellow. **r,** Quantification of angles between lateral membranes and the germline surface in AFCs and MBFCs. Mean+95% CI, two-tailed unpaired t-test, n (45 AFCs, 82 MBFCs, 4 EC). **s,** Phase field model of 3 FCs with an affinity gradient in contact with nurse cells. **k,** Illustration of cell-autonomous spreading as a function of affinity, the resulting forces and apical surface shapes in response to phase 1 and 2 affinity patterns. Grey bar represents anterior tip in egg chambers. **l,** Local z-projection of the FC junctional network. wt ECs, stained for E-Cad. Orange arrowheads point at junctions between cells of the same row that remain straight, and red arrowheads point at convex junctions between FCs of different rows with different affinities. See Supp. File S2 for detailed statistical information.

Interestingly, as Eya is also expressed in the two somatic cells that enwrap each germline cyst in testis, we asked if Eya might also control affinity-like interactions in developing spermatocytes ^39, 40^. We found that the somatic cells closely envelope each cell within the cyst and thereby maximize the soma-germline interface (Fig. S3a,c). When we expressed *eya-RNAi* in somatic cells, they failed to extend in between germline cells (Fig. S3b,c) and caused an overall change in spermatocyte morphology reflecting a minimization of the soma-germline interface (Fig. S3d). Thus, Eya controls soma-germline interfaces, possibly via regulating soma-germline affinity, in both ovaries and testis.

### Eya induces FC affinity for nurse cells in a level-dependent manner

To test if cell-cell affinity between Eya-positive FCs and nurse cells alone could account for the observed FC morphologies, we designed a phase field model that allowed us to simulate cell shapes as a function of interface dynamics ^41, 42^. We modelled 3 FCs and specified affinity as the energetic preference of an FC to be in contact with a defined boundary, representing the nurse cell surface (Fig. S3e). First, we assigned low and equal affinities to all 3 FCs, simulating low Eya levels in MBFCs during phase 2 (Fig. 3j). This gave rise to an even distribution of FCs with equal germline-contacting surfaces, recapitulating the shape of phase 2 MBFCs (Fig. 3j,k). Next, we replicated clonal ectopic expression of Eya by assigning higher affinity to the central cell (Fig. 3l, Movie 1). This caused the central cell to expand its contact with the simulated nurse cell surface at the expense of neighbouring cells, recapitulating the dominant apical surface expansion of MBFCs with ectopic Eya expression (Fig. 3m,n, Fig. S3e). Thus, translating Eya levels into differential affinities towards nurse cells was sufficient to account for Eya-dependent FC shapes.

To further explore how Eya levels determine the interaction of FCs with nurse cells, we analysed AFCs during phase 2. As described before, AFCs expand their contact surface with nurse cells in a gradient ^27^, which positively correlated with Eya levels (Fig. S3f). We found that AFCs increased their contact surface with nurse cells in a polarized manner by extending a broad actin-rich protrusion posteriorly (Fig. 3o). The polarized apical expansion deformed posterior adherens junctions, and coincided with a trailing edge-like structure on the basal side and a tilt in lateral membranes (Fig. 3o,p,q,r). To test if Eya-controlled affinity for nurse cells could give rise to such polarized cell morphologies during phase 2, we assigned a gradient of affinities based on Eya levels to the 3 simulated FCs. This recapitulated polarized protrusions of “apical” surfaces towards decreasing affinities and a tilt of lateral membranes (Fig. 3s, Movie 2). Thus, an Eya-dependent gradient in affinity for nurse cells can account for the polarized expansion of AFCs during phase 2.

Importantly, Eya-controlled affinity for nurse cells could also account for the prominent differences in FC shapes during phase 1 and 2 (Fig. 3t,u). During phase 1, uniform Eya levels in FCs translate into uniform apical expansion forces that are balanced at cell-cell junctions and thereby give rise to a regular quasi-hexagonal arrangement of cells. In contrast, an affinity gradient causes imbalanced expansion forces at junctions, which resolve into a polarized expansion towards decreasing affinities giving rise to the observed fish-scale like pattern in AFCs during phase 2. The experimental data combined with the mathematical modelling of cellular behaviors as a function of affinity propose that Eya induces level-dependent affinity of the apical FC surface for the nurse cell surface and thereby controls FC shapes.

### Eya-controlled affinity dynamics account for FC distribution over germline cells

To test if Eya-controlled affinity is sufficient to account for the shape as well as distribution of FCs throughout the 3 phases of egg chamber morphogenesis (Fig. 4a,b), we designed a more elaborate phase field model (Fig. S4a-d). We modelled 14 FCs representing 6 rows of AFCs, and for simplicity, just 5 rows of MBFCs and 3 rows of PFCs (Fig. S4b). The boundary, representing the germline surface, was divided into an affine (nurse cells) and non-affine (oocyte) compartment (Fig. S4c). Our model did not include germline growth and FC volumes were set to be constant. We measured Eya levels in rows 1-6 (AFCs) and row 7 (MBFC) in egg chambers during stages 5-10b (phase 1-3) and used the estimated length of developmental stages ^43, 44^ to interpolate the temporal development of Eya levels within each row (Fig. 4c). The resultant Eya dynamics were then used as direct proxy for affinity dynamics, with row 7 dynamics being assigned to rows 7-14 (MBFCs & PFCs). Simulating affinity based on measured Eya levels was sufficient to recapitulate FC behavior throughout development (Fig. 4d, Movie 3). During phase 1, all FCs had similar contact surface sizes, cuboidal shapes and maintained their relative positions. During phase 2, AFCs progressively increased their contact surface in a gradient from anterior to posterior and consequently displaced MBFCs (row 7-11) onto the oocyte (Fig. 4e,f). Eventually, in phase 3, FCs stably segregated with high affinity cells in contact with nurse cells and low affinity cells positioned over the oocyte. Thus, simulating FC behavior based on Eya-controlled affinity recapitulates FC positions, shapes and contact surface sizes throughout all 3 phases and is sufficient to establish a match between Eya-positive AFCs and nurse cells, as well as between Eya-negative MBFCs+PFCs and the oocyte.

**Figure 4:**
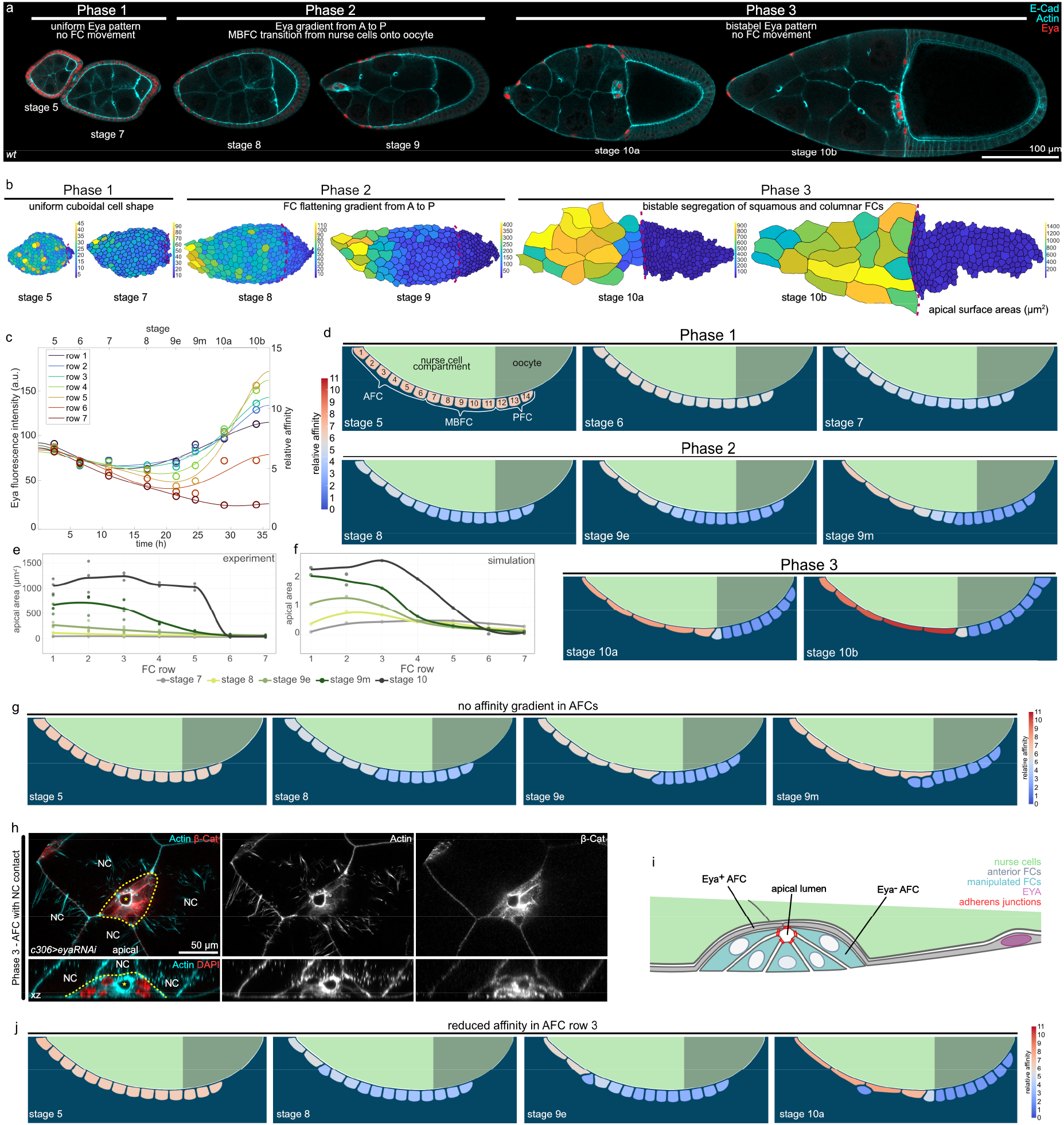
Eya-regulated affinity accounts for FC dynamics throughout egg chamber development. **a,** Medial confocal sections of ECs stained for E-Cad, F-Actin and Eya. **b,** Segmented apical surface areas of FC in ECs. Red dotted line marks nurse cell-oocyte boundary. Colour scale bar shows apical surface area sizes in (µm^2^). **c,** Average Eya fluorescence intensities in 7 anterior rows as a function of time. Time denotes hours after the beginning of stage 5. Intensities are assigned to the midpoint of each stage. Eya dynamics serve as proxy for affinity dynamics in simulations (Supp. File S1). Row 1-6 dynamics were assigned to cells 1-6 (AFCs) and row 7 dynamic was assigned to cells 7-14 (MBFCs) in the phase field model. n (stage 5: 5 EC, stage 6: 5 EC, stage 7: 5 EC, stage 8: 4 EC, stage 9e: 8 EC, stage 9m: 9 EC, stage 10a: 5 EC, stage 10b: 5 EC). 6^th^ order polynomial fit constrained to have vanishing derivatives at t = 0 and t = 36 hours. **d,** Phase field model simulating collective behaviour of FCs as a function of their affinity for germline cells. wt affinity dynamics are based on measured Eya levels. **e,** Apical surface area of 7 anterior cell rows in stage 7-10 wt ECs. Apical surface area gradient first appears at stage 8. Stage 10a and 10b were pooled (stage 10). n (stage 7: 4 EC, stage 8: 3 EC, stage 9e: 5 EC, stage 9m: 5 EC, stage 10: 3 EC) **f,** Apical surface areas of the 7 anterior cells in the simulation ((apical length)^2^). Apical surface area gradient first appears at stage 8 and develops with similar dynamics as observed *in vivo*. **g,** Phase field model simulating collective behaviour of FCs as a function of their affinity for germline cells. All 6 AFCs share row 1 high-affinity dynamics. **h,** AFCs expressing *eya-RNAi* under the control of *c306-GAL4* (patchy AFC driver, c306>*eya-RNAi*). A phase 3 EC, stained for β-cat, F-Actin and nuclei (DAPI). Formation of an apical lumen (yellow star) in *eya-RNAi* AFC cluster (yellow dotted line). NC marks nurse cells. Section through apical lumen and xz-reslice shown. **i,** Illustration of cell morphologies upon *eya-RNAi* knockdown in a group of AFCs in contact with nurse cells during phase 3. Eya-negative AFCs drastically reduce their contact surfaces with nurse cells. **j,** Phase field model simulating the collective behaviour of FCs as a function of their affinity for germline cells. Reduction in affinity in row 3 causes a failure to flatten and drives displacement of AFC from the nurse cell surface. See Supp. File S2 for detailed statistical information.

To understand if the Eya-gradient itself is required to drive proper matching of AFCs with nurse cells during phase 2, we abolished the gradient in simulations by assigning all 6 AFC rows the high-affinity dynamic of row 1 (Fig. 4g, Movie 4). This disrupted the AFC surface area gradient during phase 2, as recapitulated by experimental data (Fig. S4e,f), but more significantly, caused row 6 to displace row 7 and 8 from the nurse cell surface. This suggests that steep affinity differences between neighboring cells result in forces that are strong enough to displace low-affinity cells from the germline surface.

Lacking an experimental setup to manipulate specifically the Eya-gradient in AFCs during phase 2, we created steep differences in affinity between individual AFCs by expressing *eya-RNAi* in small AFC clones. We observed that single *eya-RNAi*-expressing AFCs as well as *eya-RNAi*-expressing AFC clones lost nurse cell contact (Fig. 4h,i, S4g,h). Single cells extruded as spheres, while clones retained epithelial features and formed a cyst with an apical lumen. When we recapitulated this experiment in simulations by reducing the affinity of cell row 3, row 3 failed to increase its contact surface area with nurse cells and was eventually displaced from the germline surface (Fig. 4j, Movie 5). Thus, steep differences in Eya levels and the resulting differences in affinities cause displacement and exclusion of low affinity FCs from nurse cells. Consequently, the gradient in Eya levels during phase 2 is essential to retain all FCs in contact with the germline while driving the redistribution of FCs to establish the right match between FCs and germline cells.

### Controlling FC distribution over the germline surface by controlling Eya-expression

If Eya levels determine whether FCs remain in contact with nurse cells or not, we should be able to control FC distribution by simply changing Eya expression patterns. To test if we can retain more FCs in contact with nurse cells, we assigned the high affinity dynamic of row 2 to rows 6-8 in our model (Fig. 5a, Movie 6). The simulation revealed an ectopic increase of their contact surfaces with nurse cells and a failure of their transition onto the oocyte, resulting in a reduced number of FCs in contact with the oocyte by phase 3. Experimentally, we tested this hypothesis by forcing MBFCs to ectopically express Eya after phase 1 using *mirr*-GAL4 (*mirr>eya*) (Fig. 5b,c). In control egg chambers, the first *mirr*-GAL4 positive cell row reached the oocyte at a germline size of 11650 µm^2^ (95% CI: 10630-12570 µm^2^) (Fig. S5a,b,c,d), defining a critical size at which ectopic *eya* expression (*eya^OE^*) by *mirr*-GAL4 was expected to prevent further FC transition onto the oocyte. As predicted by the simulation, *mirr>eya* MBFCs ectopically increased their contact surface with nurse cells and failed to transition onto the oocyte once the critical size was reached (Fig. 5d,e,f, Fig. S5e). The forced mismatch of Eya-expressing MBFCs with nurse cells caused the UMAP trajectory to divert from the control trajectory exactly at the critical germline size, highlighting the importance of appropriate germline-soma matching for overall egg chamber morphology (Fig. 5g,h, Fig. S5 f,g,h).

**Figure 5:**
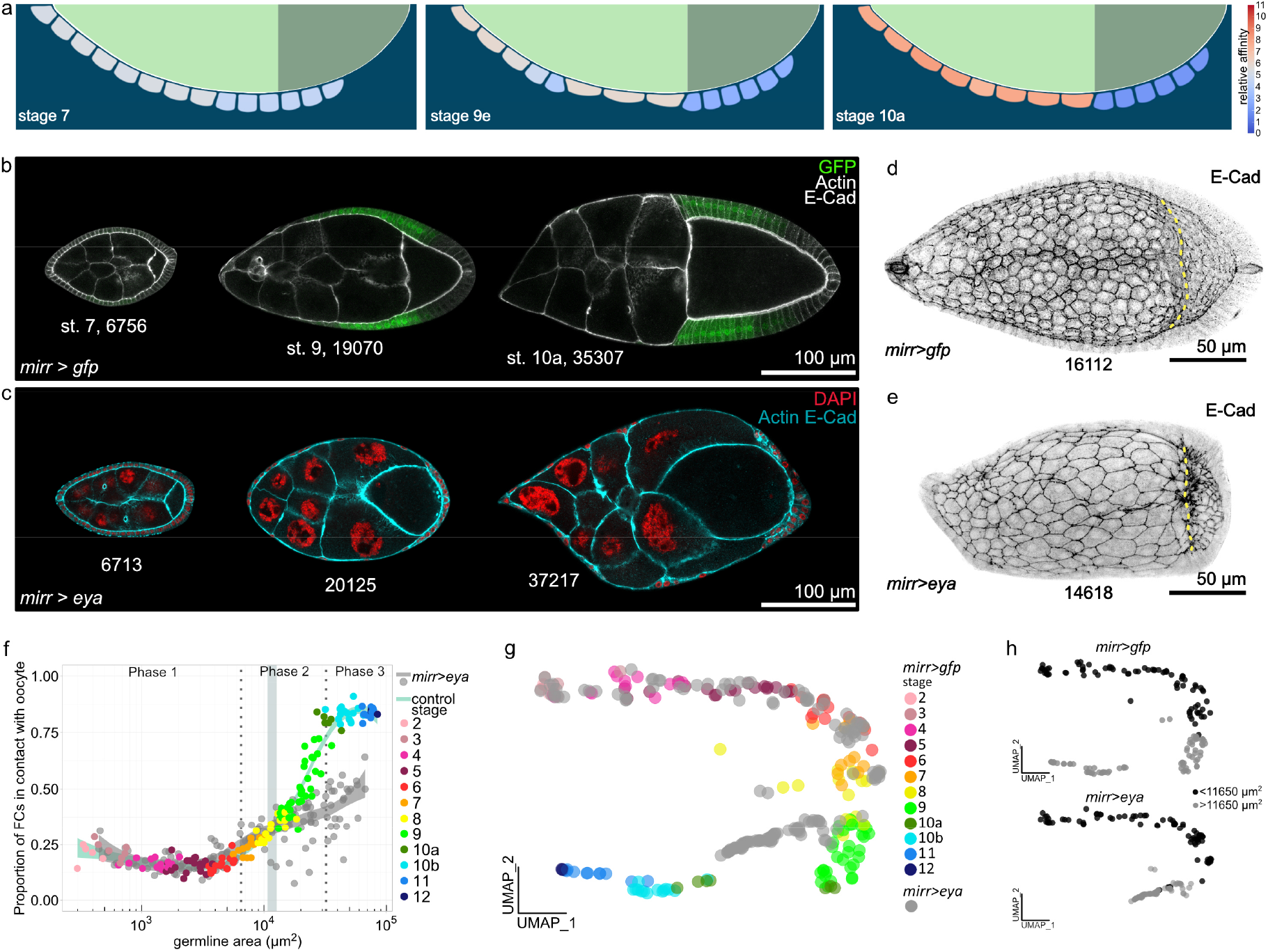
Ectopic Eya expression in MBFCs matches MBFCs with nurse cells, instead of the oocyte. **a,** Phase field model simulating the collective behaviour of FCs as a function of their affinity for germline cells. An ectopic increase of affinity in rows 6-8 was simulated. Rows ectopically increase contact with nurse cells and fail to transition onto the oocyte. **b,** Medial confocal sections of egg chambers from stage 7 to 10a expressing *gfp* under the control of *mirr-GAL4* (*mirr>gfp*, MBFC driver), stained for E-Cad and F-Actin. Note how GFP-positive cells shift from nurse cells onto the oocyte. Numbers denote germline areas in µm^2^. **c,** Medial confocal sections of egg chambers expressing *eya^OE^* under the control of *mirr-GAL4* (*mirr>eya^OE^*, MBFC driver), stained for E-Cad, F-Actin and nuclei (DAPI). Numbers denote germline areas in µm^2^. **d,** Local z-projection of the FC junctional network of egg chambers expressing *gfp* under the control of *mirr-GAL4* (*mirr>gfp*, MBFC driver), stained for E-Cad. Yellow dotted line marks oocyte-nurse cell boundary. Number denotes germline area in µm^2^. **e,** Local z-projection of the FC junctional network of egg chambers expressing *eya^OE^* under the control of *mirr-GAL4* (*mirr>eya^OE^*, MBFC driver), stained for E-Cad. Yellow dotted line marks oocyte-nurse cell boundary. Number denotes germline area in µm^2^. **f,** Proportion of FCs contacting the oocyte as a function of germline area of *mirr>gfp* and *mirr>eya^OE^* egg chambers. Bluegrey area marks 95% CI of the critical size (10632-12569 µm^2^, see Fig. S5). n (*mirr>gfp*: 153 EC; *mirr>eya^OE^*: 157 EC). **g,** UMAP plot comparing *mirr>gfp* and *mirr>eya^OE^* egg chamber morphogenesis. *mirr>eya^OE^* trajectory diverts from control trajectory during phase 2. **h,** UMAP plot of *mirr>gfp* and *mirr>eya^OE^* grouped into egg chambers with germline sizes smaller (black) or larger (grey) than the critical size (11650µm^2^, see Fig. S5). Note how the switch from black to grey in the *mirr>eya^OE^* trajectory correlates with the point of diversion from the control trajectory. See Supp. File S2 for detailed statistical information.

Thus, Eya expression patterns and resulting differential affinities dictate FC distribution over the germline. Accordingly, linking Eya expression with AFC fate ensures robust matching of AFCs with nurse cells and MBFCs+PFCs with the oocyte.

### Oocyte growth dynamics correlate with Eya-expression patterns in FCs

After characterizing how Eya expression in FCs controls their interaction with the germline, we switched perspectives and asked if Eya expression in FCs also affects how germline cells interact with FCs. To analyse a possible differential interaction of nurse cells and the oocyte with the follicle epithelium, we characterized the angle at the interface, where oocyte and nurse cells compete for FC contact (Fig. 6a). The angle depicts which germline cell type preferentially expands its contact surface with the follicle epithelium, and thereby serves as a read-out for whether nurse cells or the oocyte harbour effective affinity for the follicle epithelium (Fig. 6b). An angle of 90° is the result of balanced forces, whereas an angle >90° represents effective nurse cell affinity and an angle <90° effective oocyte affinity. We found that the angle was larger than 90° during phase 1, decreased below 90° during phase 2, and increased above 90° again in phase 3 (Fig. 6c,d). This suggested that the oocyte harbours effective affinity for FCs exclusively during phase 2. We then analysed Eya levels in FCs overlying the nurse cell-oocyte boundary and found that these appeared with similar dynamics as the interface angle (Fig. 6e). To characterize the relationship between Eya levels and the interface angle, we performed a linear regression of the interface angle as a function of Eya levels and found a positive correlation, with Eya levels <72 a.u. predicting an interface angle <90° (effective oocyte affinity) and Eya levels >72 a.u. predicting an interface angle >90° (effective nurse cell affinity) (Fig. 6f). A phase-wise analysis suggested that exclusively during phase 2 Eya levels in FCs at the nurse cell-oocyte boundary are low enough to establish effective oocyte affinity (Fig. 6g). In line with that, we found that the contact surface of the oocyte with the follicle epithelium only started to increase at the transition from phase 1 to phase 2 and came to a halt at the end of phase 2, before it further increased during nurse cell dumping in phase 3 (Fig. 6h). Remarkably, the increase in interface with the follicle epithelium was accompanied by an increase in oocyte size (Fig. 6i).

**Figure 6:**
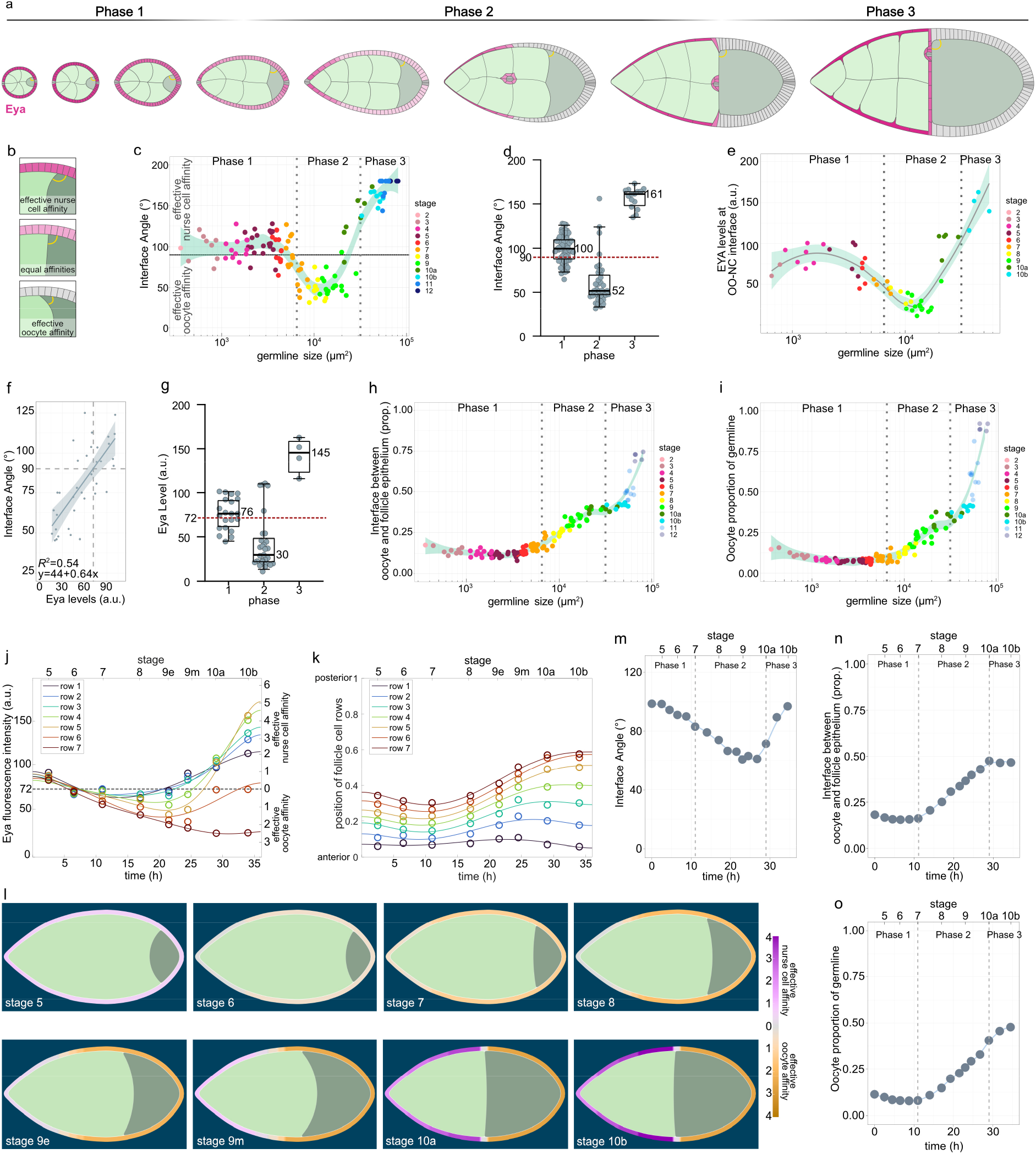
Eya-controlled effective germline affinities account for oocyte growth and shape dynamics. **a,** Illustrations of ECs depicting the interface angle and Eya expression patterns. **b,** Illustration of the interface angle as a parameter characterizing effective affinities of oocyte or nurse cell towards FCs . **c,** Interface angle as a function of the germline area. Dotted lines mark germline sizes at the transition between two phases. LOESS fitted with a 95% CI area. n=126 EC. **d,** Interface angle of wt ECs grouped into the three phases. Box plot with whiskers marking the 5^th^ and 95^th^ percentile. Numbers state the median of the corresponding phase. n (phase 1: 62 EC, phase 2: 39 EC, phase 3: 15 EC) **e,** Eya levels in FCs at the nurse cell-oocyte boundary as a function of germline area. Dotted lines mark germline sizes at the transition between two phases. LOESS fitted with a 95% CI area. n=54 EC. **f,** Linear regression between the interface angle and Eya levels in FCs in contact with the nurse cell-oocyte boundary. Dashed line marks 90° angle. Linear regression with 95% CI area, n= 36 EC. **g,** Eya levels in FCs at nurse cell-oocyte boundary of wt ECs grouped into the three phases. Box plot with whiskers marking the 5^th^ and 95^th^ percentile. Number states median of the corresponding phase. n (phase 1: 19 EC, phase 2: 29 EC, phase 3: 4 EC) **h,** Oocyte-FC interface proportion of germline-FC interface as a function of germline area. LOESS fitted with a 95% CI area. n=126 EC. **i,** Oocyte area proportion of the germline as a function of the germline area. Dotted lines mark germline sizes at the transition between two phases. LOESS fitted with a 95% CI area. n=126 EC. **j,** Average Eya fluorescence intensity of the first 7 anterior FC rows as a function of time. Time denotes hours after the beginning of stage 5. Average Eya intensities are assigned to the midpoint of each stage. Eya dynamics in rows serve as proxy for effective affinity dynamics implemented in simulations (Supp. File S1). n (stage 5: 5 EC, stage 6: 5 EC, stage 7: 5 EC, stage 8: 4 EC, stage 9e: 8 EC, stage 9m: 9 EC, stage 10a: 5 EC, stage 10b: 5 EC). 6^th^ order polynomial fit constrained to have vanishing derivatives at t = 0 and t = 36 hours. **k,** Average normalized distances to anterior pole of the first 7 anterior FC rows as a function of time. Distances and intensity dynamics were used to simulate effective affinities of germline cells (Supp. File S1). Row 7 affinity dynamic was assigned from the distance of row 7 to the posterior pole. n (stage 5: 5 EC, stage 6: 5 EC, stage 7: 5 EC, stage 8: 4 EC, stage 9e: 8 EC, stage 9m: 9 EC, stage 10a: 5 EC, stage 10b: 5 EC). **l,** Phase field model simulating germline cell behaviour as a function of their affinity for the follicle epithelium. wt affinity dynamics are based on measured Eya levels. **m,n,o** Individual morphological parameters of the simulation as a function of time. **m,** Interface Angle. **n,** Proportion of the germline-FC interface made up by the oocyte. **o,** Oocyte proportion of the germline. Phase boundaries were assigned to mid stage 7 and mid stage 10a. See Supp. File S2 for detailed statistical information.

### Eya-controlled effective germline affinities account for oocyte growth and shape dynamics

We therefore hypothesized that Eya levels in FCs non-cell autonomously regulate the contact surfaces of germline cells with the follicle epithelium and that this subsequently controls oocyte expansion. To test this hypothesis, we designed a phase field model, simulating the oocyte and the nurse cell compartment, and defined the outer boundary as the contact surface with FCs (Fig. S6a,b). We quantified Eya levels in FC rows from stage 5-10b and the row’s relative position along the AP-axis, covering all three morphogenetic phases (Fig. 6j,k). We used these spatio-temporal dynamics as proxy for effective-affinity-inducing conditions on the boundary, with Eya levels of 72 a.u. inducing 0 effective affinity (Fig. 6j). The resulting simulation recapitulated interface angle and oocyte expansion dynamics very well (Fig. 6l, Movie 7). The interface angle was >90° during phase 1 and dropped below 90° at the end of phase 1, when the switch from effective nurse cell to effective oocyte affinity took place (Fig. 6m). This switch also caused an expansion of the oocyte contact surface with FCs and oocyte growth (Fig. 6n,o). Once, the oocyte had expanded along the entire Eya-negative FC surface, the angle increased above 90° and the oocyte seized to grow. Thus, these simulations suggest that Eya in FCs controls effective germline affinity for the follicle epithelium, causing the oocyte to expand its surface specifically along Eya-negative FCs. Next to Eya-controlled FC redistributions, this would result in an additional morphogenetic dynamic ensuring the establishment of the right match between FCs and germline cells.

### Premature loss of Eya in FCs during phase 1 induces premature oocyte expansion

To test the role of FC Eya expression in controlling oocyte expansion, we first manipulated Eya expression in phase 1. We hypothesized that the uniform expression of Eya above the critical level of 72 a.u. during phase 1 resulted in effective nurse cell affinity, which prevented the oocyte from increasing its contact with FCs and grow in size. In line with that, simulating an early switch to effective oocyte affinity during phase 1 led to a premature decrease in the interface angle below 90° and a premature expansion of the oocyte (Fig. 7a-d, Movie 8). Experimentally, we reduced Eya levels in phase 1 egg chambers by GR1-GAL4 driven *eya-RNAi* (*gr1>eya-RNAi*) (Fig. 7e,f), which led to a substantial decrease in Eya levels when egg chambers reached a germline size of 1600 µm^2^ (95% CI: 1300-1918 µm^2^) (Fig. S7a,b). The loss of Eya caused a premature decrease of the interface angel below 90°, indicating a switch from effective nurse cell affinity to effective oocyte affinity (Fig. 7g, Fig. S7c). The premature loss of effective nurse cell affinity also caused a loss of nurse cell organization within the germline cyst (Fig. 7h). In control egg chambers, nurse cells were of similar sizes and distributed evenly along the follicle epithelium, whereas a loss of Eya in FCs resulted in a high variance of nurse cell-FC interface lengths and nurse cell sizes (Fig. 7i,j, Fig. S7d,e). Thus, Eya expression in FCs non-autonomously controls nurse cell arrangement and morphology. Furthermore, as predicted by simulations, the *eya-RNAi*-driven switch from effective nurse cell affinity to effective oocyte affinity caused a premature expansion of the oocyte-FC interface and oocyte size (Fig. 7k,l, Fig. S7f,g,h). The disruption of the *gr1>eya-RNAi* UMAP trajectory from the control during phase 1 further highlighted the importance of FC Eya expression and the resulting soma-germline affinity for global egg chamber morphogenesis (Fig. 7m,n). Taken together, during phase 1, uniform Eya levels in FCs above the critical level give rise to effective nurse cell affinity, which inhibits premature oocyte expansion and is essential to organize the germline cyst.

**Figure 7:**
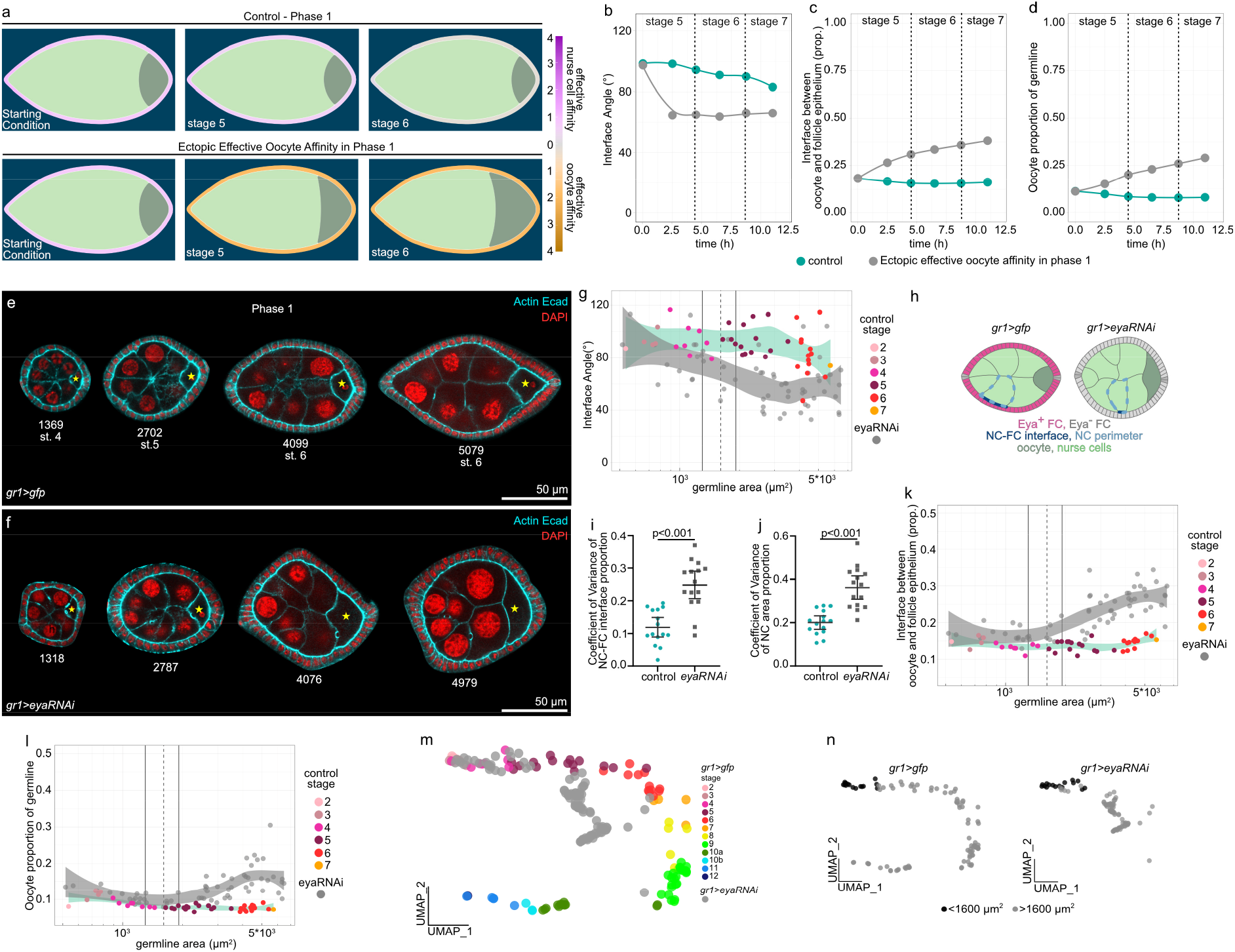
Premature loss of Eya in FCs during phase 1 induces premature oocyte expansion. **a,** Phase field model simulating germline cell behaviour as a function of their affinity for the follicle epithelium. wt affinity vs. ectopic effective oocyte affinity in phase 1 (simulating phase 1 Eya loss). **b,c,d** Individual morphological parameters of the simulation as a function of time for phase 1. **b,** Interface Angle. **c,** Proportion of the germline-FC interface made up by the oocyte. **d,** Oocyte proportion of the germline. **e, f** Medial confocal sections of phase 1 ECs expressing *gfp* (e) or *eya-RNAi* (f) under the control of *gr1-GAL4* (*gr1>gfp*, *gr1>eya-RNAi*, FC driver), stained for E-Cad, F-Actin and nuclei (DAPI). Yellow stars mark oocytes. Numbers denote germline areas in µm^2^. **g,** Interface Angle as a function of germline area of *gr1>gfp* and *gr1>eya-RNAi* ECs in phase 1 (germline area <6500µm^2^). Dotted line marks critical germline area and solid lines mark 95% CI. All curves are LOESS fitted with a 95% CI area. n (*gr1>gfp* = 42 EC, *gr1>eya-RNAi* = 60 EC). **h,** Illustrations of *gr1>gfp* and *gr1>eya-RNAi* egg chambers in phase 1 (stage 6). **i,** Quantification of the coefficient of variance (CV) of the nurse cell-FC interface (dark blue line) proportion of the nurse cell perimeter (light blue line) within a nurse cell cluster. Mean+95%CI, two-tailed unpaired Student’s t-test, n (*gr1>gfp*: 15 EC, 71 NCs; *gr1>eya-RNAi*: 15 EC, 89 NCs). **j,** Quantification of the coefficient of variance (CV) of nurse cell area proportions within a nurse cell cluster. Mean+95%CI, two-tailed unpaired Welch’s t-test, n (*gr1>gfp*: 15 EC, 71 NCs; *gr1>eya-RNAi*: 15 EC, 89 NCs). **k,** Oocyte-FC interface proportion of germline-FC interface as a function of germline area of of *gr1>gfp* and *gr1>eya-RNAi* ECs in phase 1 (germline area <6500µm^2^). **l,** Oocyte area proportion of the germline area as a function of germline area of *gr1>gfp* and *gr1>eya-RNAi* ECs in phase 1 (germline area <6500µm^2^). Dotted line marks critical germline area and solid lines mark 95% CI of critical germline area. Curves are LOESS fitted with a 95% CI area. n (*gr1>gfp* = 42 EC, *gr1>eya-RNAi* = 60 EC). **m,** UMAP plot comparing *gr1>gfp* and *gr1>eya-RNAi* EC morphogenesis. **n,** UMAP plots of *gr1>gfp* and *gr1>eya-RNAi* coloured by germline sizes smaller (black) and larger (grey) than the critical size (1600 µm^2^, see Fig. S6). See Supp. File S2 for detailed statistical information.

### Ectopic Eya expression in MBFCs inhibits oocyte expansion

Next, we hypothesized that the loss of Eya in MBFCs and the resultant switch to effective oocyte affinity drives oocyte expansion during phase 2. To test this, we simulated ectopic effective nurse cell affinity in the region of MBFCs after phase 1 (Fig. 8a, Movie 9). As a consequence, the interface angle failed to decrease, and the oocyte failed to expand its contact with FCs and to increase in size during phase 2 (Fig. 8b-d). Experimentally, we forced MBFCs to ectopically express Eya during phase 2 using *mirr*-GAL4 (*mirr>eya*) (Fig. 8e,f). As predicted, ectopic Eya expression in MBFCs retained the interface angle above 90° representing a failure in switching to effective oocyte affinity (Fig. 8g, Fig. S8a), once egg chambers had reached the critical size for mirr-GAL4-driven interference (Fig. S5a,b,c,d). Accordingly, the oocyte failed to increase its interface with FCs, as posterior nurse cells outcompeted the oocyte for FC contact disrupting germline cluster organization (Fig. 8h,i). Ultimately, the ectopic effective nurse cell affinity resulted in a failure of oocyte growth (Fig. 8j, Fig. S8b,c). Thus, the loss of Eya expression in MBFCs and the resultant switch to effective oocyte affinity is essential for oocyte growth and consequently egg chamber morphogenesis (Fig. 5g,h, Fig. S5 f,g,h).

**Figure 8:**
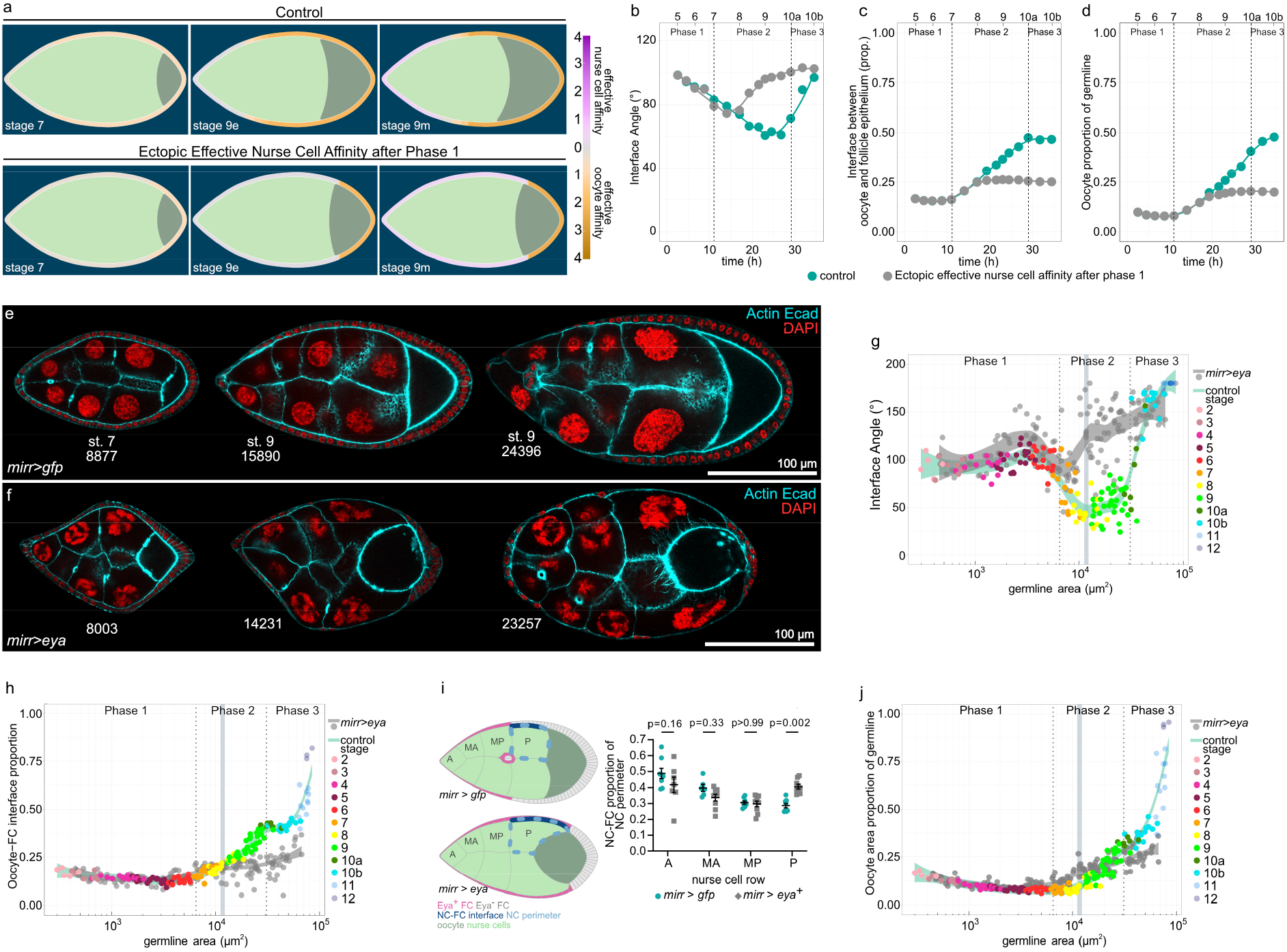
Ectopic Eya expression in MBFCs induces ectopic effective nurse cell affinity and inhibits oocyte growth. **a,** Phase field model simulating germline cell behaviour as a function of their affinity for the follicle epithelium. wt affinity dynamics vs. ectopic effective nurse cell affinity after phase 1 (simulating Eya expression in MBFC after phase 1). **b,c,d** Individual morphological parameters of the simulation as a function of time in phase 1. **b,** Interface Angle. **c,** Proportion of the germline-FC interface made up by the oocyte. **d,** Oocyte proportion of the germline. Phase boundaries were assigned to mid stage 7 and mid stage 10a. **e, f** Medial confocal sections of ECs expressing *gfp* (e) or *eya* (f) under the control of *mirr-GAL4* (*mirr>gfp*, *mirr>eya*^OE^, MBFC driver), stained for E-Cad, F-Actin and nuclei (DAPI). Numbers denote germline areas in µm^2^. **g,h** Individual morphological parameters as a function of germline area for *mirr>gfp* and *mirr>eya^OE^* ECs. **g,** Interface Angle. **h,** Proportion of the germline-FC interface made up by the oocyte. Dotted lines mark germline sizes at the transition between two phases. Bluegrey area marks 95% CI of the critical size (10632-12569 µm^2^). All curves are LOESS fitted with a 95% CI area. n (*mirr>gfp*: 153 EC; *mirr>eya*^OE^: 157 EC). **i,** Illustrations depicting phase 2 *mirr>gfp* and *mirr>eya^OE^* egg chambers and the quantification of the nurse cell perimeter (light blue line) proportion made up by the nurse cell-FC interface (dark blue line) for individual nurse cells in *mirr>gfp* and *mirr>eya^OE^* ECs. Nurse cells are grouped by nurse cell row (A=anterior, MA=mid-anterior, MP=mid-posterior, P=posterior). Nurse cell row averages of ECs were analysed. Mean±SE, two-way Anova with Šídák’s multiple comparisons test, n (*mirr>gfp* (A: 8 EC, MA: 6 EC, MP: 8 EC, P: 8 EC), *mirr>eya^OE^* (A: 7 EC, MA: 9 EC, MP: 9 EC, P: 9 EC)). **j,** Oocyte proportion of the germline as a function of germline area for *mirr>gfp* and *mirr>eya^OE^* ECs. Dotted lines mark germline sizes at the transition between two phases. Bluegrey area marks 95% CI of the critical size (10632-12569 µm^2^). All curves are LOESS fitted with a 95% CI area. n (*mirr>gfp*: 153 EC; *mirr>eya*^OE^: 157 EC). See Supp. File S2 for detailed statistical information.

### Creating an entirely Eya-negative follicle epithelium results in ectopic oocyte expansion

We found that establishing the right match between germline cells and FCs by the end of phase 2 correlated with a switch back to effective nurse cell affinity and a cessation of oocyte expansion up until nurse cell dumping is initiated mid phase 3 (Fig. 6c-g). We hypothesized that the match of the oocyte with all available Eya-negative FCs and the consequent positioning of Eya-positive FCs at the nurse cell-oocyte boundary at the transition from phase 2 to 3 causes the temporary halt in oocyte expansion. We thus tested if we could override this halt in oocyte growth by turning the entire epithelium Eya-negative. We simulated this by modelling effective oocyte affinity along the entire interface from phase 2 onwards. This caused the interface angle to remain smaller than 90° and resulted in a continuation of oocyte expansion after phase 2 (Fig. 9a-d, Movie 10). *In vivo*, we expressed a constitutively active EGFR under the control of TJ-Gal4 (*tj>egfr^λtop^*) throughout the epithelium, which prohibited FCs at the anterior tip to adopt AFC fate ^20^, consequently producing egg chambers with an Eya-negative follicle epithelium from phase 2 onwards (Fig. 9e,f). We found that these egg chambers retained their interface angles below 90° even after phase 2, reflecting prolonged effective oocyte affinity (Fig. 9g, Fig. S8d). In line with that, the Eya-negative follicle epithelium failed to halt oocyte expansion at the end of phase 2 (Fig. 9h,i, Fig. S8e,f), which caused a disruption of egg chamber morphogenesis (Fig. 9j,k, Fig. S8g,h). Hence, the oocyte expands its contact exclusively with Eya-negative FCs and therefore halts once the right match is established at the end of phase 2. Thus, our data demonstrate that Eya in FCs non-cell autonomously controls the interaction of germline cells with the follicle epithelium, and thereby regulates oocyte growth dynamics. Taken together, Eya in FCs drives the matching of FCs and germline cells by controlling soma-germline coordination in a bilateral manner, giving rise to a robust matching mechanism (Fig. 10).

**Figure 9:**
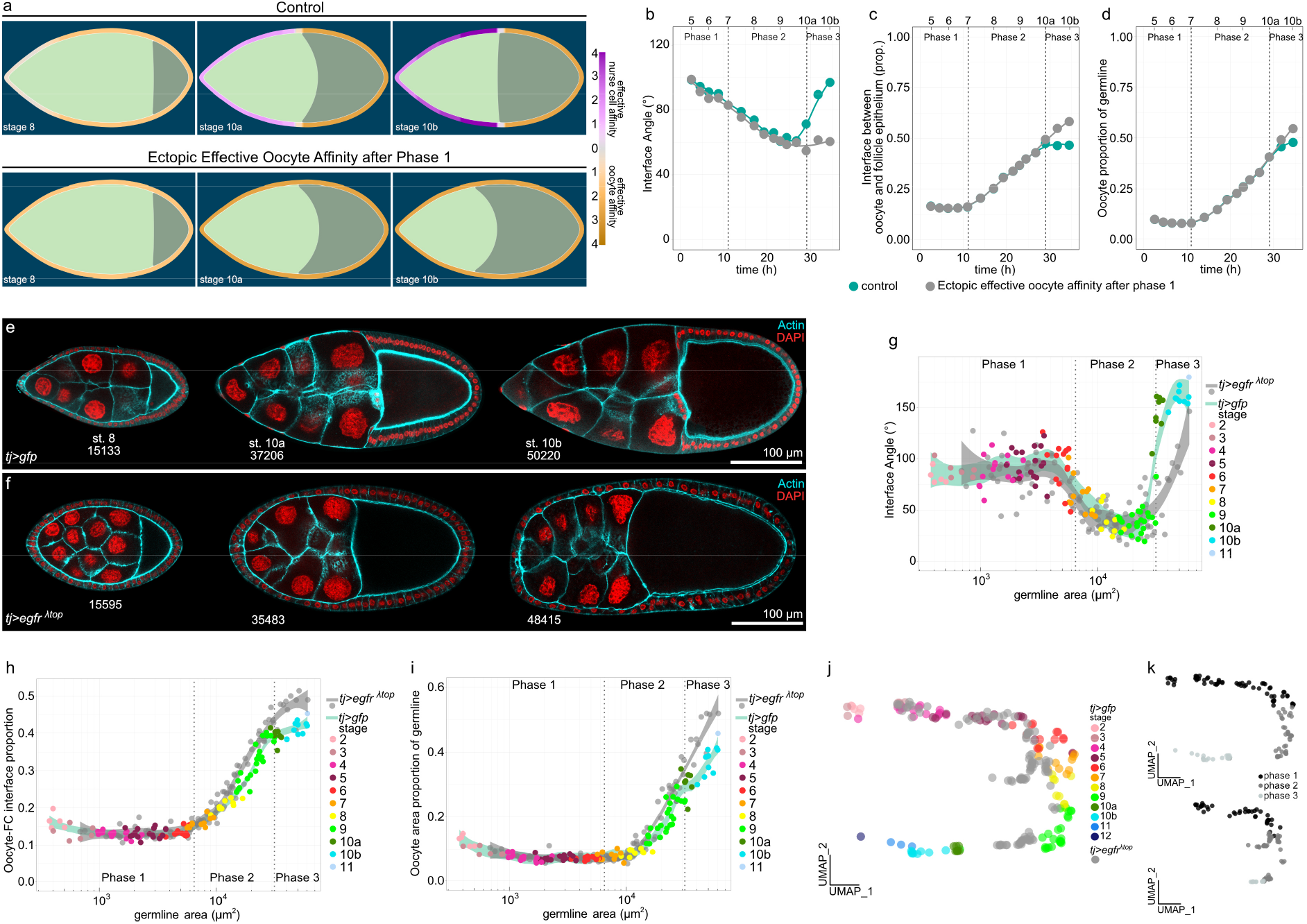
Inhibition of AFC differentiation causes ectopic oocyte affinity and oocyte expansion during phase 3. **a,** Phase field model simulating germline cell behaviour as a function of their affinity for the follicle epithelium. wt affinity dynamics vs. ectopic effective oocyte affinity after phase 1 (simulating an Eya-negative follicle epithelium after phase 1). **b,c,d** Individual morphological parameters of the simulation as a function of time for phase 1. **b,** Interface Angle. **c,** Proportion of the germline-FC interface made up by the oocyte.**d,** Oocyte proportion of the germline. Phase boundaries were assigned to mid stage 7 and mid stage 10a. **e,** Medial confocal sections of egg chambers expressing *CD8*-tom and gfp under the control of *tj-GAL4* (*tj>gfp*, FC driver), stained for F-Actin and nuclei (DAPI). Numbers denote germline areas in µm^2^. **f,** Medial confocal sections of egg chambers expressing *CD8*-*tom* and *egfr*^λtop^ under the control of *tj-GAL4* (*tj>egfr*^λtop^, FC driver), stained for F-Actin and nuclei (DAPI). Numbers denote germline areas in µm^2^. **g,h,i** Individual morphological parameters as a function of germline area for *tj>gfp* and tj> *egfr*^λtop^ egg chambers. **g,** Interface Angle. **h,** Proportion of the germline-FC interface made up by the oocyte. **i,** Oocyte proportion of the germline. Dotted lines mark germline sizes at the transition between two phases. Curves are LOESS fitted with a 95% CI area. n (*tj>gfp*: 119 EC, *tj>egfr*^λtop^: 109 EC). **j,** UMAP plot comparing *tj>gfp* and *tj>egfr*^λtop^ egg chamber morphogenesis. **k,** UMAP plots of *tj>gfp* and *tj>egfr*^λtop^ grouped into phase 1 (black, germline area<6500µm^2^), phase 2 (darkgrey, germline area>6500µm^2^ & <31500µm^2^), and phase 3 egg chambers (lightgrey, germline area>31500µm^2^). See Supp. File S2 for detailed statistical information.

**Figure 10:**
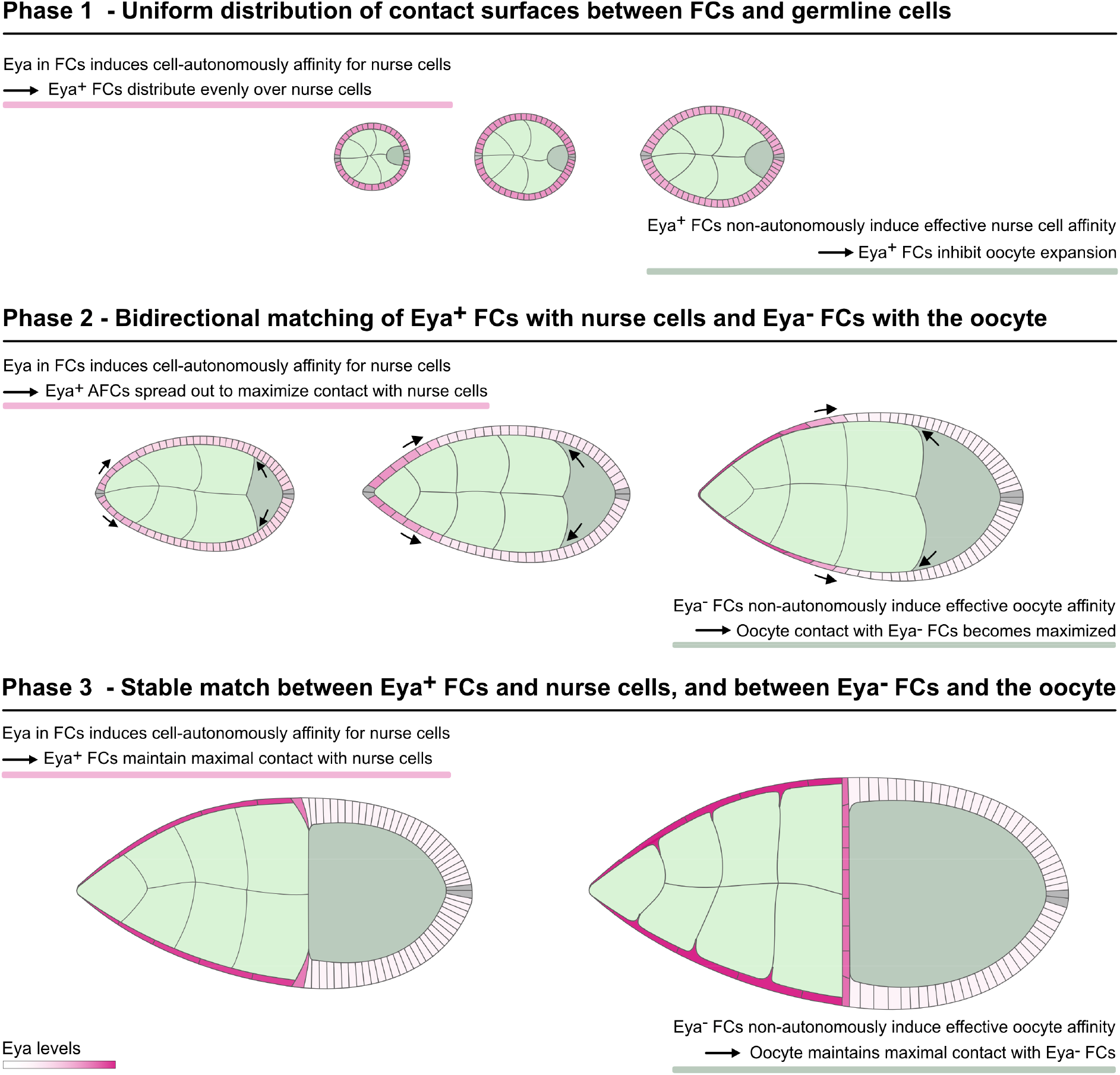
Eya-driven matching of FCs and germline cells. Illustration of Eya-controlled soma-germline coordination during *Drosophila* egg chamber morphogenesis. Eya in FCs cell-autonomously induces affinity for nurse cells, but not for the oocyte. This causes Eya-positive AFCs to spread out posteriorly which maximizes their contact with nurse cells, and consequently displaces Eya-negative MBFCs onto the oocyte. Additionally, Eya in FCs non-autonomously controls effective germline affinity, such that Eya-positive FCs induce effective nurse cell affinity, while Eya-negative FCs induce effective oocyte affinity. As a result, the oocyte expands anteriorly during phase 2 and maximizes its contact exclusively with Eya-negative FCs. Ultimately, these bilateral affinity dynamics result in bidirectional matching dynamics that ensure robust matching of Eya-positive FCs with nurse cells and Eya-negative FCs with the oocyte. The established match represents the energetically preferred and therefore highly stable state.

## Discussion

Our study demonstrates how germline and soma self-organize into functional units by using differential cell-cell affinity to match cell populations across cell lineages. We identify the co-transcriptional regulator Eya as the master regulator of this process. Our data demonstrate that Eya-expressing FCs cell-autonomously experience affinity towards nurse cells, but not towards the oocyte, while Eya-negative FCs experience affinity for neither. Additionally, we show that Eya in FCs regulates non-cell-autonomously the interaction of germline cells with the follicle epithelium, such that nurse cells experience effective affinity for Eya-positive FCs, whereas the oocyte experiences effective affinity for Eya-negative FCs. Moreover, our experiments demonstrate that these Eya-controlled bilateral affinity dynamics underlie the critical matching of FC subpopulations with germline cells.

Importantly, our phase field simulations provide a controlled environment to isolate the effects of changing affinity between cell lineages. We see that an affinity differential between cell types proportional to their relative Eya expression very closely recovers experimentally observed morphologies of both FCs and the germline. Furthermore, with parameters only adapted to wild-type models, the numerical simulations correctly predict significant morphological changes, such as FC-germline contact loss or premature oocyte expansion when affinities are suitably altered. This lends additional credence to our finding that Eya controlled differential affinity of cell lineages is underlying soma-germline matching.

Taken together, we demonstrate that Eya-controlled bilateral affinity dynamics at the soma-germline interface create a robust self-organizing system to drive the establishment of inter-lineage functional units. Thus, it represents a prime example of how differential cell-cell affinity can be utilized to drive complex morphogenetic events of multi-lineage tissues.

Yet, molecular and cellular details of Eya-dependent interactions remain unknown. A previous study revealed that MBFCs display high levels of apical-medial myosin when in contact with nurse cells and that MBFCs lose apical-medial myosin as soon as they encounter the oocyte ^22^. While we show that Eya downregulation in MBFCs is sufficient to remove affinity for nurse cells and displace MBFCs onto the oocyte, the contact-dependent myosin enrichment suggests that Eya-negative FCs may not just lack affinity for nurse cells but experience active repulsion from the nurse cell surface. However, either mechanism will ensure reliable matching between Eya-negative MBFCs and the oocyte.

Furthermore, as we cannot experimentally separate the interaction of nurse cell or the oocyte with FCs, we introduced ‘effective affinities’ to characterize the interaction of germline cells with FCs in a relative manner. Consequently, we cannot distinguish whether effective oocyte affinity is the result of active cell-autonomous affinity of the oocyte for Eya-negative FCs or the consequence of a repulsion between nurse cells and Eya-negative FCs. However, both scenarios will result in the same effective affinity which would control oocyte expansion.

The most pressing question might be how differential affinity at the soma-germline interface is established at the molecular level. We expect that Eya regulates the expression of transmembrane receptors in FCs, which recognize nurse cell or oocyte-specific ligands or receptors. Subsequently, signaling downstream of these receptors must alter interfacial tensions by targeting the cytoskeleton and adhesion complexes in FCs and germline cells ^2, 3^.

## Author Contributions

Conceptualization VW, PD, AKC

Validation VW, PD, AKC

Investigation VW, PD, AKC

Writing VW, AKC

Visualization VW, PD, AKC

Supervision AKC

Funding Acquisition AKC

## Acknowledgements

We thank the LIC facility at the University of Freiburg for technical help with imaging. We thank the Bloomington Drosophila Stock Center (BDSC), the Vienna Drosophila Stock Collection (VDRC) and the Developmental Studies Hybridoma Bank (DSHB) for providing fly stocks and antibodies. The finite element simulations of the phase field model were performed using the zec C++ framework developed by L. Courte and M. Zeinhofer. We thank David Bilder, Martin Zeidler, Vincent Mirouse, Sally Horne-Badovinac, Kim McCall, Thomas Lecuit, Ulrich Tepass, Barry Thompson and Giorgos Pyrowolakis for sharing reagents. We thank Lance Fredrick Pahutan Bosch and Katrin Kierdorf for help and discussion regarding the UMAP analysis. We thank Peter Walentek for valuable comments on the manuscript. The authors acknowledge support by the state of Baden-Württemberg through bwHPC.

---

**Supplemental Figure S1:**
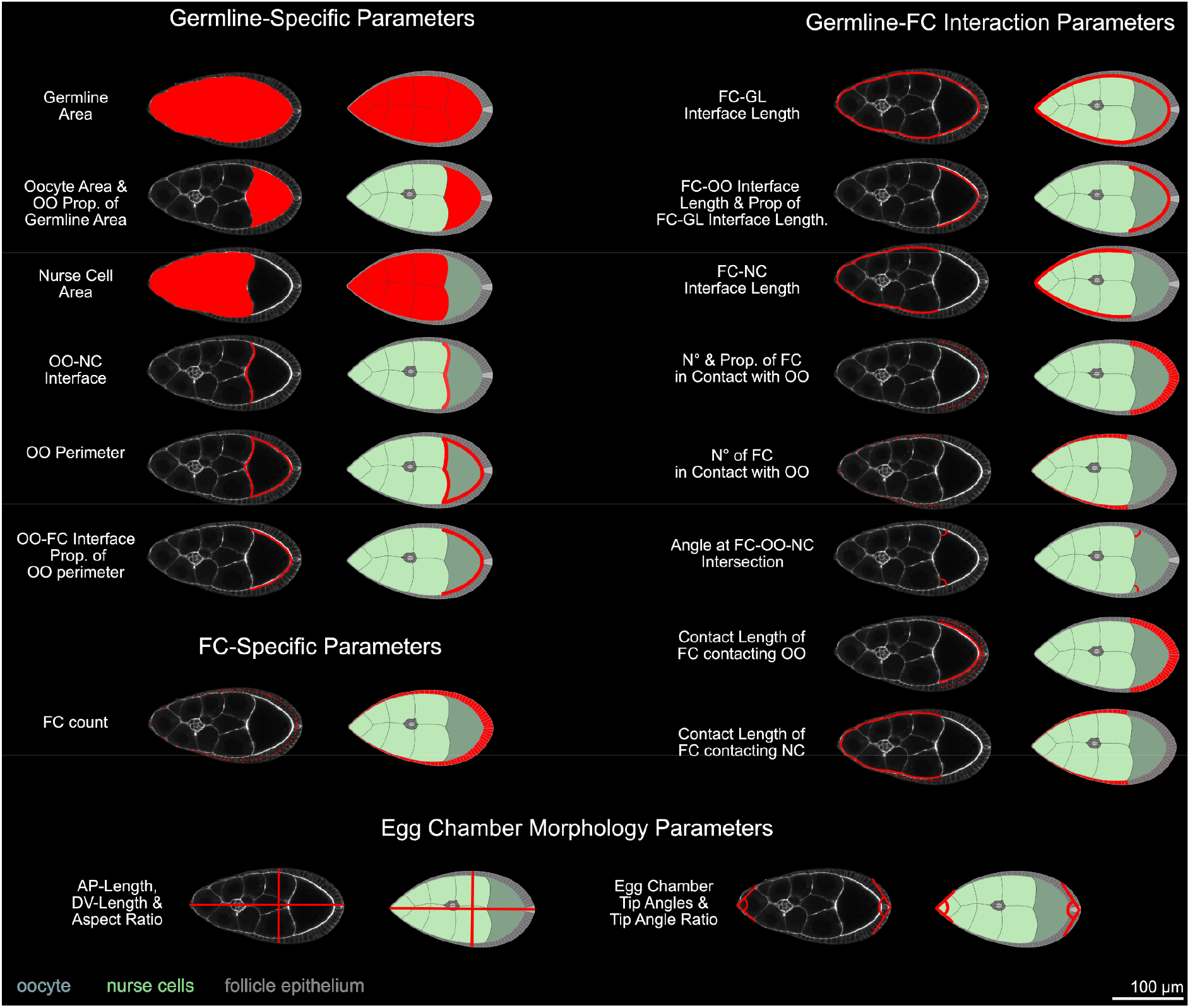
Drosophila egg chamber morphology parameters. Medial confocal sections and illustrations of phase 2 egg chambers visualizing parameters (red) that were quantified for the multidimensional morphology description.

**Supplemental Figure S2:**
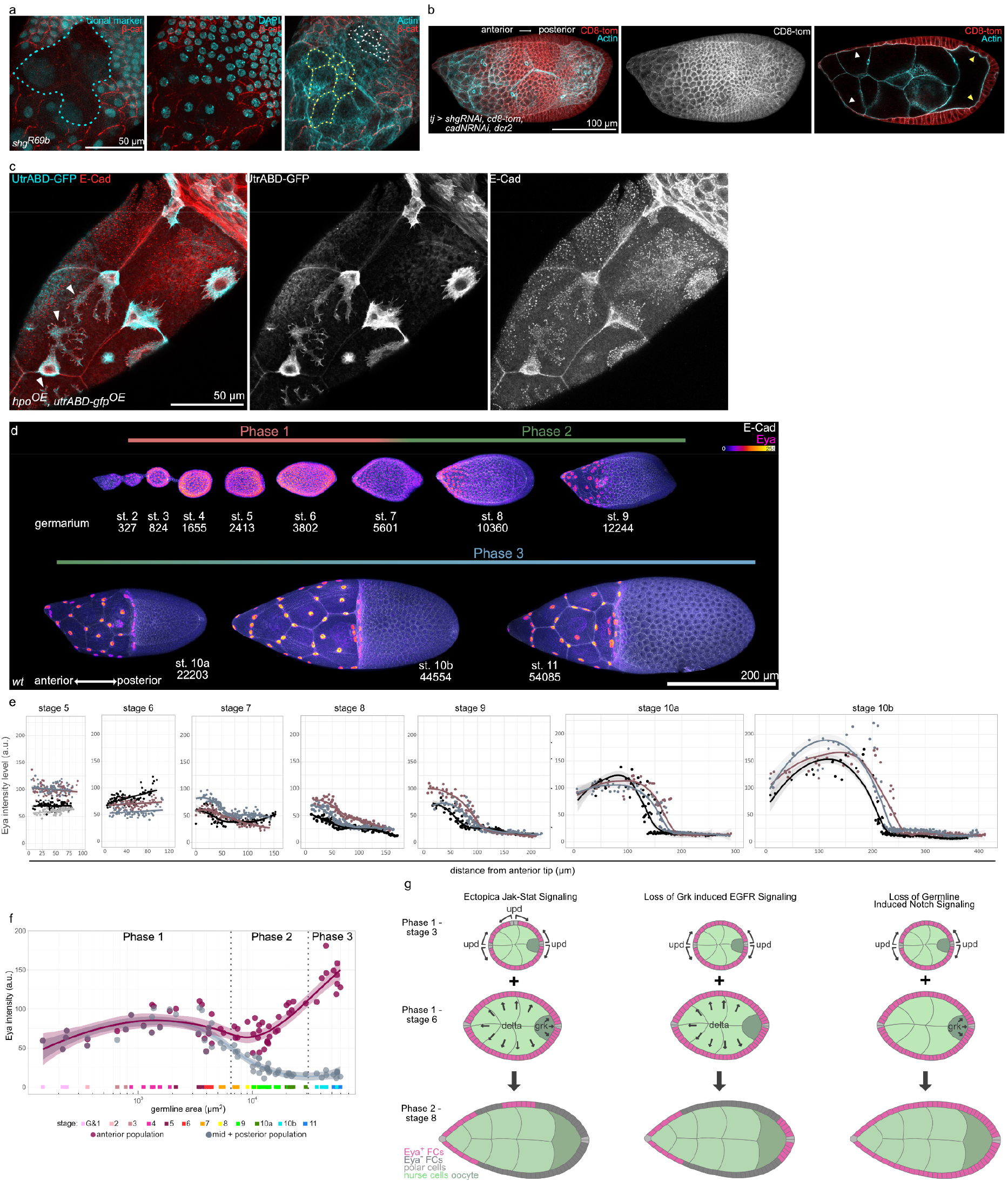
AFCs spread actively over the nurse cell surface. **a,** Maximum fluorescence intensity projection of a *shg^R69b^* (null mutant) clone (cyan dotted outline) in a phase 2 (stage 9) egg chamber consisting of AFCs and MBFCs. Stained for β-cat, F-Actin and nuclei (DAPI). Apical surfaces of AFCs (yellow) and MBFCs (white) are outlined. E-Cad loss does not disrupt AFC flattening, and MBFCs maintain their comparatively small apical areas. **b,** Maximum fluorescence intensity projection of an egg chamber expressing s*hg-RNAi, cadN-RNAi, CD8-tom* and *dcr2* under the control of *tj-GAL4* (FC driver), stained for F-Actin. Loss of E-Cad and N-Cad causes disruption of PFC morphology (yellow arrowheads), but not of AFC spreading (white arrowheads). **c,** Maximum fluorescence intensity projection of a phase 3 egg chamber with clonal overexpression of *hippo (hpo)* and *utrABD-gfp*, stained for E-Cad. *hpo* overexpression leads to reduced cell volume. AFCs detach from each other but continue to spread out cell autonomously (white arrowheads point at protrusions). **d,** Maximum fluorescence intensity projections of representative egg chambers covering all three morphological phases, stained for E-Cad and Eya. Fire LUT visualizes Eya levels in nuclei of FCs throughout egg chamber development. Numbers denote germline area in µm^2^. **e,** Nuclear Eya levels in FCs as a function of their distance to the anterior pole of egg chambers from stages 5 to 10b. Colours represent individual egg chambers at each stage. Colours do not relate between stages. n (st. 5: 4 EC, st. 6-10b: 3 EC). Curves are LOESS fitted with a 95% CI area. **f,** Measured mean Eya intensities in anterior (maroon) and mid+posterior (grey) FC populations of egg chambers as a function of germline area. Coloured squares represent developmental stages of egg chambers. Curves are LOESS fitted with a 95% CI area. (n=66 egg chambers). See Supp. File S2 for detailed stage-wise statistical comparison. **g,** Illustrations of genetic manipulations targeting fate determining factors and the result on Eya expression in FCs of phase 2.

**Supplemental Figure S3:**
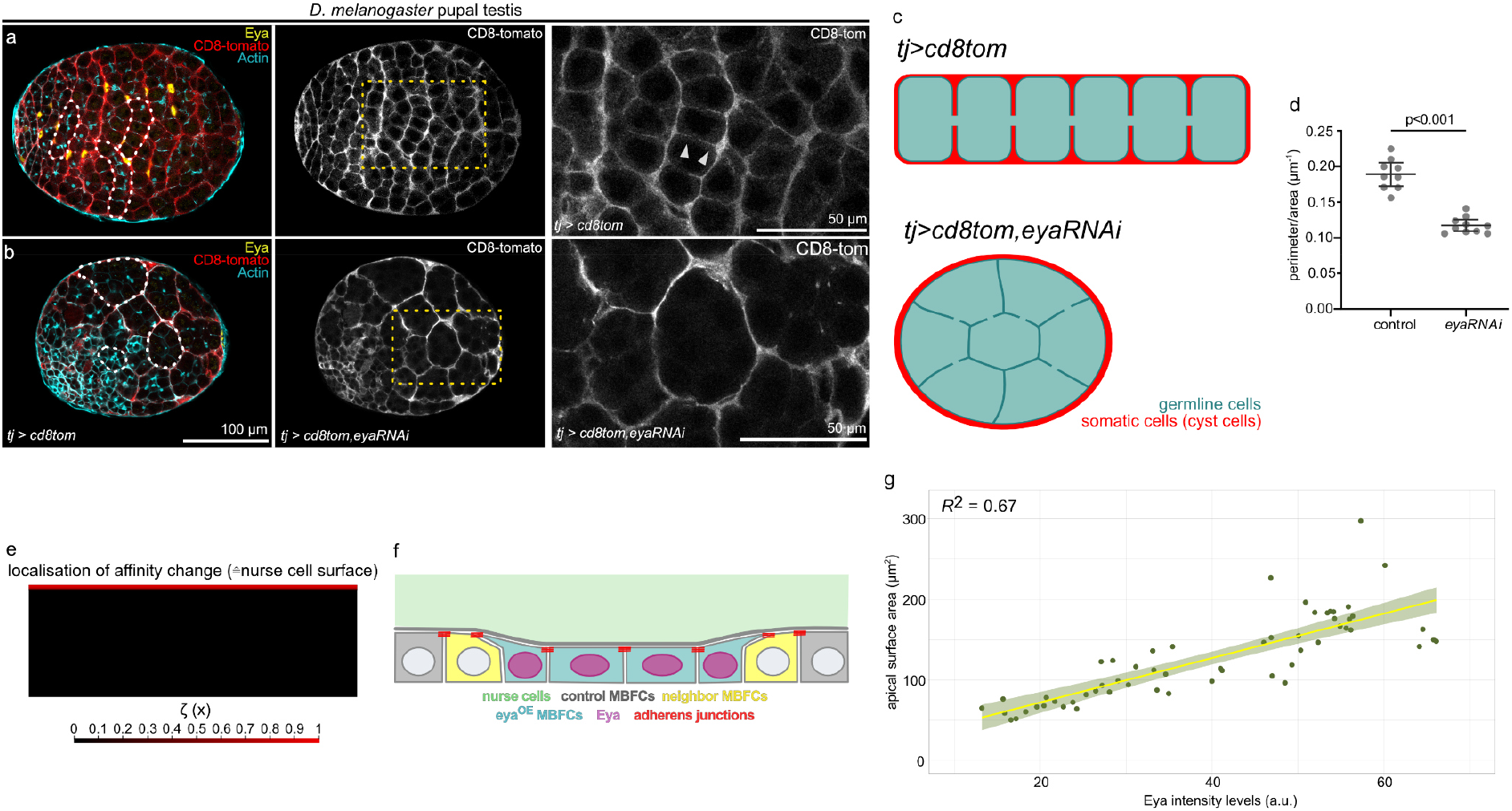
Maximization of the soma-germline interface in testis depends on Eya. **a,** Confocal sections of *D. melanogaster* pupal testis expressing *CD8-tom* under the control of *tj-GAL4* (*tj>CD8tom*, cyst cell driver). *CD8-tom* visualizes somatic cells (cyst cells) that envelope the developing germline. Somatic cells extend between individual germline cells and thereby maximize the soma-germline contact surface (white arrowheads). White dotted lines mark individual germline cysts. Yellow dotted rectangular marks position of enlarged area. **b,** Confocal sections of *D. melanogaster* pupal testis expressing *CD8-tom* and *eya-RNAi* under the control of *tj-GAL4* (*tj>CD8tom,eya-RNAi,* cyst cell driver). *CD8-tom* visualizes somatic cells (cyst cells) that envelope the developing germline. Note that loss of Eya causes failure of cyst cells extending between individual germline cells and that the germline cyst adopts a spherical shape. Consequently, the contact surface between somatic cells and germline cells is minimized. White dotted lines mark individual germline cysts. Yellow dotted rectangular marks position of enlarged area. **c,** Illustrations of *tj>CD8-tom* and *tj>CD8-tom,eya-RNAi* spermatogonial cysts. **d,** Quantification of the ratio between the germline cyst interface in contact with somatic cells and germline area. Mean+95%CI, two-tailed unpaired Student’s t-test, n (*tj>CD8-tom*: 9 cysts, *tj>CD8-tom,eya-RNAi*: 10 cysts). **e,** Localization of the affinity change that represents the nurse cell surface. **f,** Illustration of cell morphologies upon ectopic *eya^OE^* expression in MBFC clones in contact with nurse cells during phase 2. Corresponds to Fig. 3m. **g,** Linear regression between apical surface areas and Eya levels of FCs (FC from anterior rows 1-7 of stage 9 egg chambers, phase 2). Linear regression with 95% CI area, n= 57 AFCs from 3 EC. See Supp. File S2 for detailed statistical information.

**Supplemental Figure S4:**
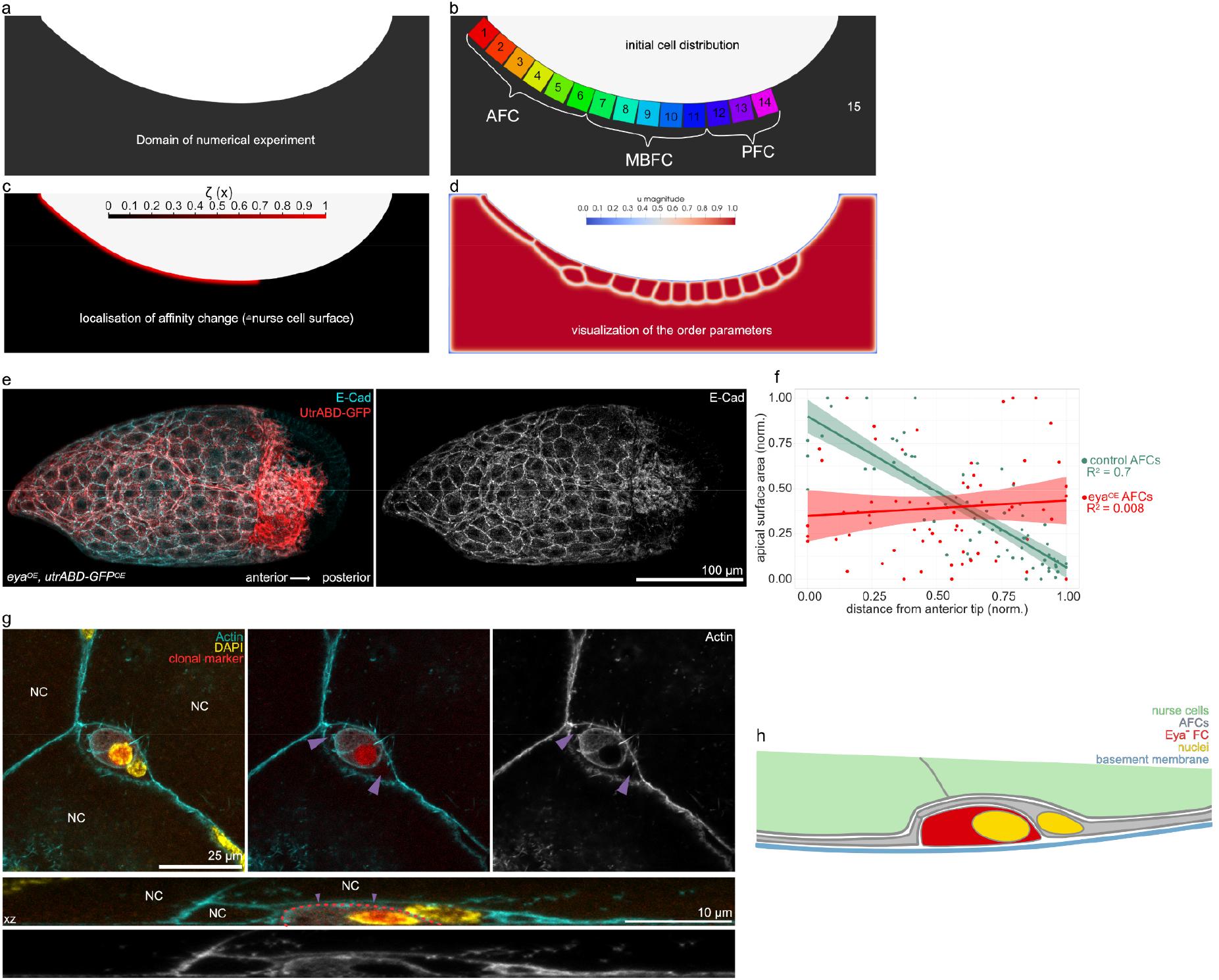
Affinity-controlled FC interaction with the germline drives FC shapes and movements. **a,** Domain of computation Ω of the numerical experiment for the phase field model. **b,** Initial cell distribution in the phase field model. Cells numbered from anterior to posterior. **c,** Localization of the affinity change that represents the nurse cell surface. **d,** Visualization of the order parameters of the phase field simulation. **e,** Maximum fluorescence intensity projection of an egg chamber with clonal expression of *eya^OE^* and *utrABD-gfp*, stained for E-Cad. Note how the gradient in apical surface areas of AFCs is lost upon broad overexpression of *eya^OE^*. **f,** Quantification of apical areas of AFCs as a function of their distance to the anterior tip for control AFCs and *eya^OE^* AFCs. Linear regression with 95% CI area. n (control FCs: 71 FC, 3 EC; *eya^OE^* FCs: 65 FC, 3 EC) **g,** AFCs with clonal *eya-RNAi* expression of a phase 3 egg chamber, stained for F-Actin and nuclei (DAPI). One AFC is expressing *eya-RNAi* (red). *Eya-RNAi* expressing AFC (red dotted line) is rounded up and disconnected from nurse cells (NC). Wild type AFC extends between *eya-RNAi* AFC and nurse cells (purple arrowheads). Confocal section and xz-reslice shown. **h,** Illustration of an AFC lacking Eya in a phase 3 egg chamber. The Eya-negative AFC is displaced from the nurse cell surface by wild type AFCs.

**Supplemental Figure S5:**
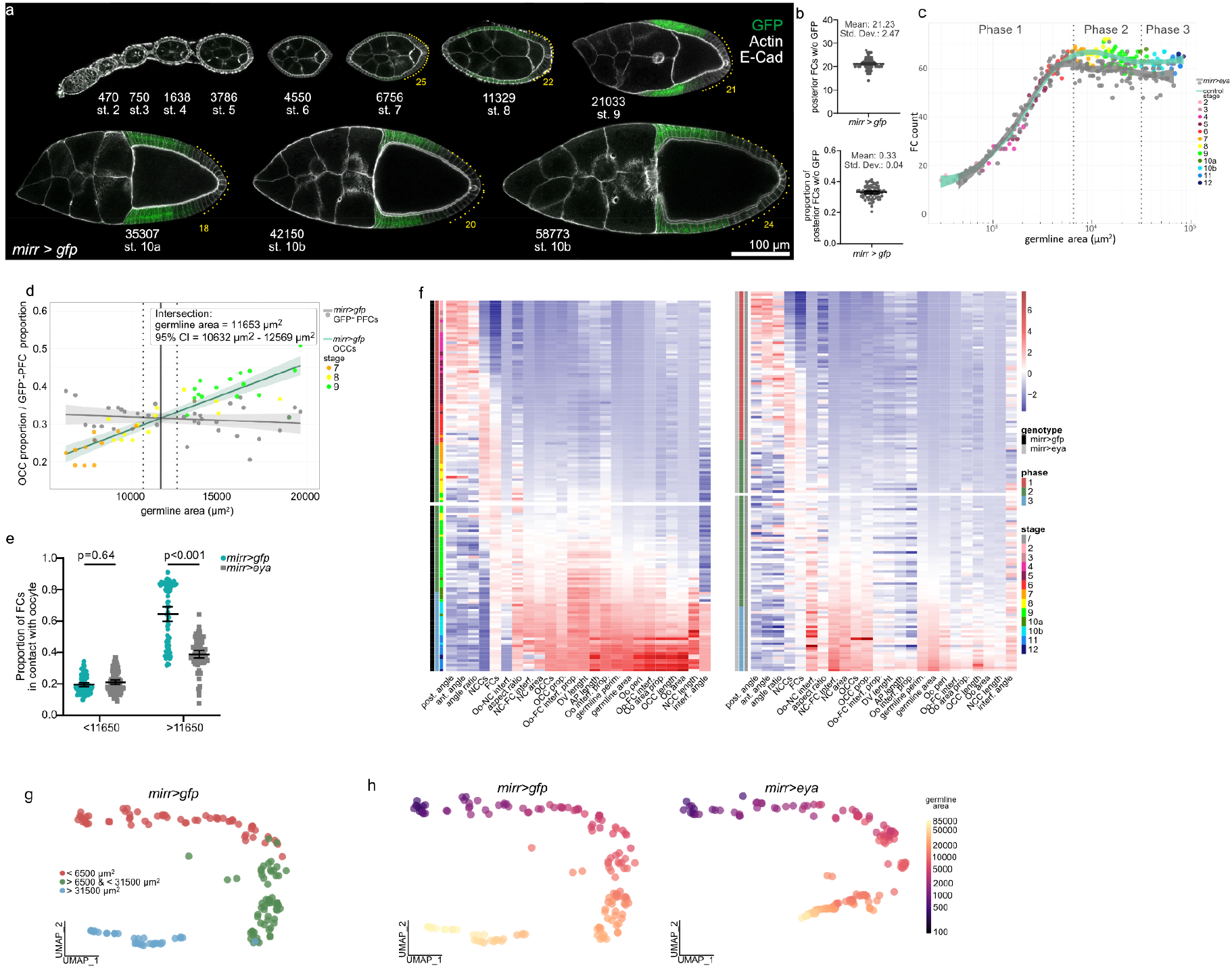
Ectopic Eya expression in MBFCs during phase 2 inhibits MBFC transition onto the oocyte. **a,** Medial confocal sections of egg chambers expressing *gfp* under the control of *mirr-GAL4* (*mirr>gfp*, MBFC driver), stained for F-Actin and E-Cad. Yellow dots mark posterior cells that are not under the control of *mirr-GAL4*. Numbers denote germline areas in µm^2^. **b,** Quantification of posterior cells without GFP as total cell count and as proportion of all FCs. Mean+95% CI, n = 80 EC. **c,** FC count as a function of germline area for *mirr>gfp* and *mirr>eya^OE^* egg chambers. LOESS fitted curves with 95% CI area. **d,** Determining the ‘critical size’ as the germline area at which first GFP-positive FCs are expected to come into contact with the oocyte. *mirr>gfp* egg chambers with germline areas > 6500 µm^2^ and < 20000 µm^2^ were used. Linear regression between the proportion of FCs in contact with the oocyte (OCC=oocyte contacting FCs) and the proportion of posterior FCs without GFP (GFP-negative PFCs) was performed. Crossing point of the two linear regression curves was fitted. Linear regression +95% CI area shown. Solid grey line marks germline area at estimated intersection point and dotted grey lines mark 95% CI of the intersection germline area. See Supp. File S2 for detailed statistical information. **e,** Parameter comparison between *mirr>gfp* and *mirr>eya* egg chambers grouped by germline area into smaller and larger than critical size (11650 µm^2^). Mean +95% CI, two-way Anova with Šídák’s multiple comparisons test; n (*mirr>gfp* (<11650 µm^2^): 74 EC, *mirr>gfp* (>11650 µm^2^): 74 EC, *mirr> eya^OE^* (<11650 µm^2^): 86 EC, *mirr> eya^OE^* (>11650 µm^2^): 71 EC). **f,** Heatmap of the 24 morphological parameters of *mirr>gfp* and *mirr>eya^OE^* egg chambers. Each row represents an individual egg chamber with increasing germline areas from top to bottom. Break in heatmap marks critical size (11650 µm^2^). n (*mirr>gfp*: 153 egg chambers, *mirr>eya*^OE^: 157 egg chambers), **g,** UMAP plot depicting *mirr>gfp* egg chambers coloured based on their phase affiliation. **h,** UMAP plot of *mirr>gfp* and *mirr>eya^OE^* egg chambers coloured by their germline area. See Supp. File S2 for detailed statistical information.

**Supplemental Figure S6:**
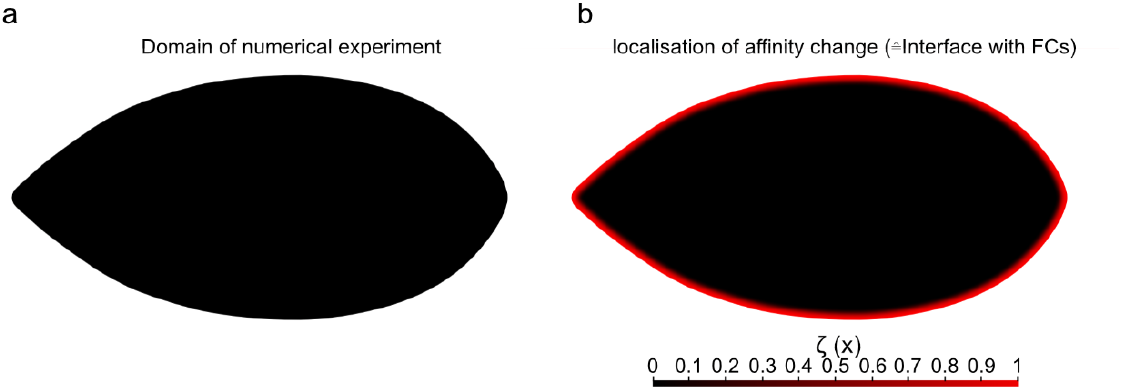
Phase Field Model of Germline Cell Behaviour as a Function of their Effective Affinity for the Follicle Epithelium. **a,** Domain of computation Ω of the numerical experiment for the phase field model. **b,** Localization of the affinity change that represents the nurse cell surface.

**Supplemental Figure S7:**
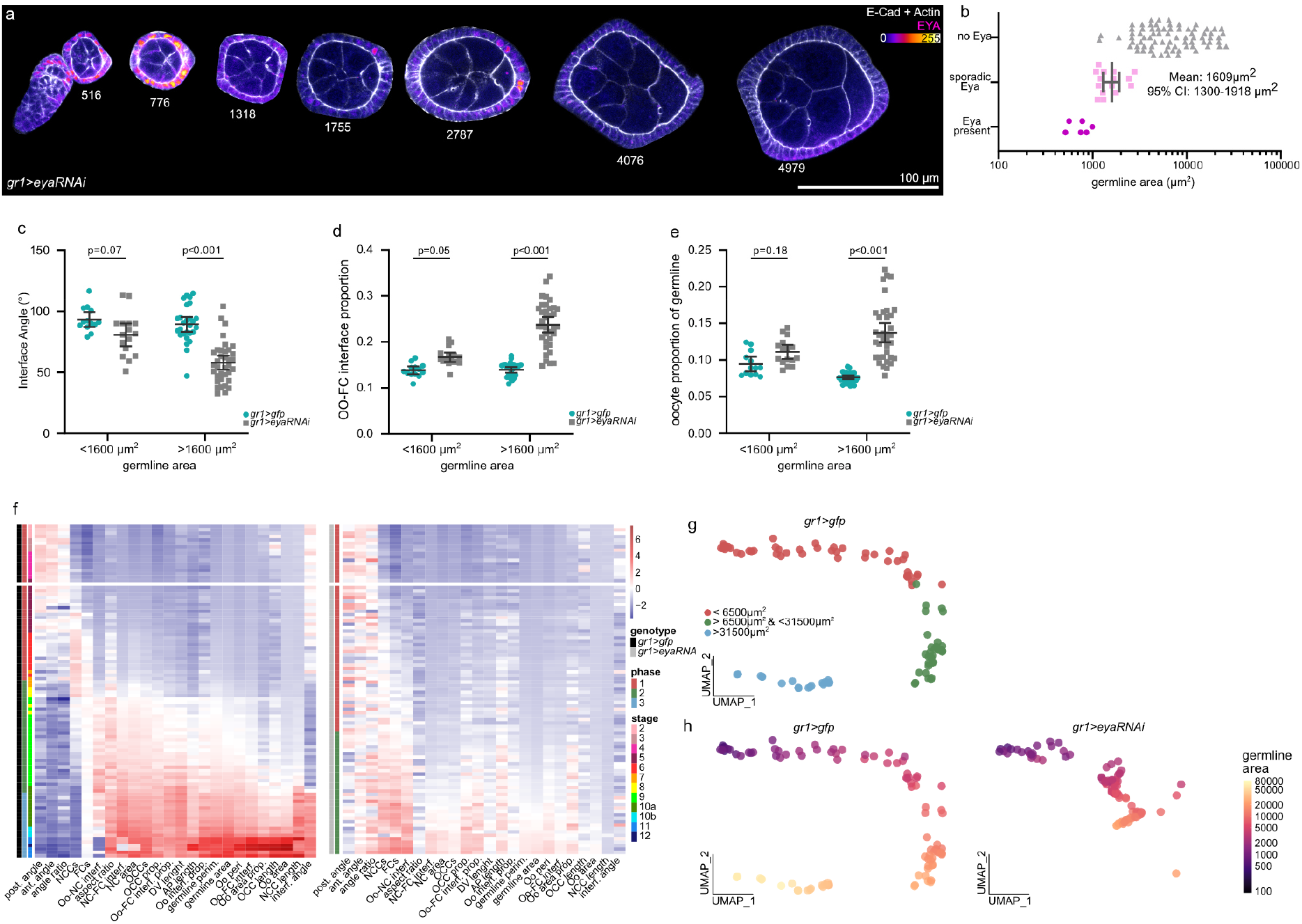
Premature loss of Eya during phase 1 disrupts egg chamber morphogenesis. **a,** Medial confocal sections of egg chambers expressing *eya-RNAi* under the control of *gr1-GAL4* (*gr1>eya-RNAi*, FC driver), stained for F-Actin, E-Cad and Eya. Numbers denote germline area. **b,** Quantification of Eya expression in FCs. Egg chambers were grouped into three categories (Eya present, sporadic Eya and no Eya in FCs) and plotted against their germline area. Mean+95%CI of the sporadic Eya group was determined as critical size from which on effects of *eya-RNAi* expression could be expected. Mean+95% CI, n (Eya present: 6 EC, sporadic Eya: 15 EC, no Eya: 75 EC). **c,d,e,** Parameter comparison between *gr1>gfp* and *gr1>eya-RNAi* egg chambers for phase 1 (germline area < 6500 µm^2^). Egg chambers grouped into smaller and larger than critical germline area (1600 µm^2^). **c,** Interface Angle. **d,** Oocyte-FC interface proportion of germline-FC interface. **e,** Oocyte area proportion of germline area. Mean+95% CI, two-way Anova with Šídák’s multiple comparisons test; n (*gr1>gfp* (<1600 µm^2^): 13 EC, *gr1>gfp* (>1600 µm^2^): 27 EC, *gr1>eya-RNAi* (<1600 µm^2^): 16 EC*, gr1>eya-RNAi* (>1600 µm^2^): 36 EC). **f,** Heatmap of the 24 morphological parameters of *gr1>gfp* and *gr1>eya-RNAi* egg chambers. Each row represents an individual egg chamber with increasing germline areas from top to bottom. Note that no egg chambers of *gr1>eya-RNAi* exist with germline sizes corresponding to phase 3, as they degenerate before. Break in heatmap marks critical size (1600 µm^2^). n (*gr1>gfp*: 97 EC, *gr1>eya-RNAi*: 96 EC). **g,** UMAP plot of *gr1>gfp* egg chambers coloured based on their phase affiliation. **h,** UMAP plot of *gr1>gfp* and *gr1>eya-RNAi* egg chambers coloured by their germline area size. See Supp. File S2 for detailed statistical information.

**Supplemental Figure S8:**
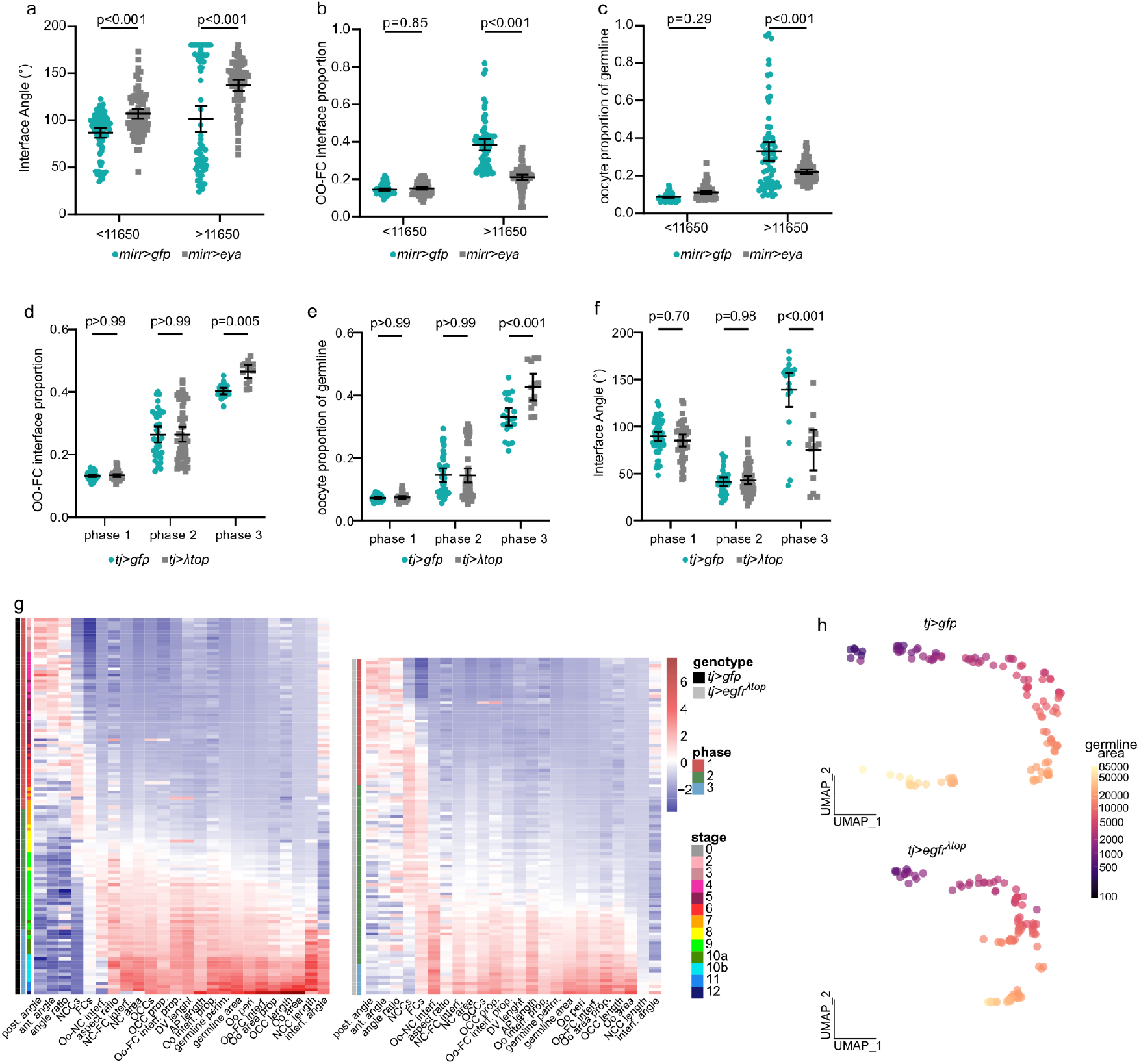
Manipulating Eya expression patterns during phase 2 and 3 disrupts oocyte expansion dynamics. **a,b,c,** Parameter comparison between *mirr>gfp* and *mirr>eya^OE^* egg chambers. Egg chambers grouped into smaller and larger than critical germline area (11650 µm^2^). **a,** Interface Angle. **b,** Oocyte-FC interface proportion of germline-FC interface. **c,** Oocyte area proportion of germline area. Mean+95% CI, two-way Anova with Šídák’s multiple comparisons test; n (*mirr>gfp* (<11650 µm^2^): 74 EC, *mirr>gfp* (>11650 µm^2^): 74 EC, *mirr>eya^OE^* (<11650 µm^2^): 86 EC, *mirr>eya^OE^* (>11650 µm^2^): 71 EC). **d,e,f** Parameter comparison between *tj>gfp* and *tj>egfr*^λtop^ egg chambers. Egg chambers subdivided by phases. **d,** Interface Angle. **e,** Oocyte-FC interface proportion of germline-FC interface. **f,** Oocyte area proportion of germline area. Mean+95% CI, two-way Anova with Šídák’s multiple comparisons test; n (*tj>gfp* (phase 1): 62 EC, *tj>gfp* (phase 2): 39 EC, *tj>gfp* (phase 3): 21 EC, *tj>egfr*^λtop^ (phase 1): 41 EC, *tj>egfr*^λtop^ (phase 2): 58 EC, *tj>egfr*^λtop^ (phase 3): 10 EC). See Supp. File S2 for detailed statistical information. **g,** Heatmap of the 24 morphological parameters of *tj>gfp* and *tj>egfr*^λtop^ egg chambers. Each row represents an individual egg chamber with increasing germline areas from top to bottom. n (*tj>gfp*: 122 EC, *tj>egfr*^λtop^: 109 EC). **h,** UMAP plot of *tj>gfp* and *tj>egfr*^λtop^ egg chambers coloured based on germline area size. See Supp. File S2 for detailed statistical information.

## References

1. Xavier da Silveira Dos Santos, A. & Liberali, P. From single cells to tissue self-organization. FEBS J 286, 1495–1513, doi:10.1111/febs.14694 (2019).

2. Gomez-Galvez, P., Anbari, S., Escudero, L. M. & Buceta, J. Mechanics and self-organization in tissue development. Semin Cell Dev Biol 120, 147–159, doi:10.1016/j.semcdb.2021.07.003 (2021).

3. Tsai, T. Y., Garner, R. M. & Megason, S. G. Adhesion-Based Self-Organization in Tissue Patterning. Annu Rev Cell Dev Biol, doi:10.1146/annurev-cellbio-120420-100215 (2022).

4. Doherty, C. A., Amargant, F., Shvartsman, S. Y., Duncan, F. E. & Gavis, E. R. Bidirectional communication in oogenesis: a dynamic conversation in mice and Drosophila. Trends in cell biology 32, 311–323, doi:10.1016/j.tcb.2021.11.005 (2022).

5. Rodrigues, P. et al. Germ-Somatic Cell Interactions Are Involved in Establishing the Follicle Reserve in Mammals. Front Cell Dev Biol 9, 674137, doi:10.3389/fcell.2021.674137 (2021).

6. Lei, L. & Spradling, A. C. Mouse oocytes differentiate through organelle enrichment from sister cyst germ cells. Science 352, 95–99, doi:10.1126/science.aad2156 (2016).

7. Niu, W. & Spradling, A. C. Mouse oocytes develop in cysts with the help of nurse cells. Cell 185, 2576–2590 e2512, doi:10.1016/j.cell.2022.05.001 (2022).

8. Hinnant, T. D., Merkle, J. A. & Ables, E. T. Coordinating Proliferation, Polarity, and Cell Fate in the Drosophila Female Germline. Front Cell Dev Biol 8, 19, doi:10.3389/fcell.2020.00019 (2020).

9. Lebo, D. P. V. & McCall, K. Murder on the Ovarian Express: A Tale of Non-Autonomous Cell Death in the Drosophila Ovary. Cells 10, doi:10.3390/cells10061454 (2021).

10. Duhart, J. C., Parsons, T. T. & Raftery, L. A. The repertoire of epithelial morphogenesis on display: Progressive elaboration of Drosophila egg structure. Mech Dev 148, 18–39, doi:10.1016/j.mod.2017.04.002 (2017).

11. Clarke, H. J. Regulation of germ cell development by intercellular signaling in the mammalian ovarian follicle. Wiley Interdiscip Rev Dev Biol 7, doi:10.1002/wdev.294 (2018).

12. Horne-Badovinac, S. & Bilder, D. Mass transit: epithelial morphogenesis in the Drosophila egg chamber. Developmental dynamics : an official publication of the American Association of Anatomists 232, 559–574, doi:10.1002/dvdy.20286 (2005).

13. Bai, J. & Montell, D. Eyes absent, a key repressor of polar cell fate during Drosophila oogenesis. Development 129, 5377–5388, doi:10.1242/dev.00115 (2002).

14. Deng, W. M., Althauser, C. & Ruohola-Baker, H. Notch-Delta signaling induces a transition from mitotic cell cycle to endocycle in Drosophila follicle cells. Development 128, 4737–4746 (2001).

15. Lopez-Schier, H. & St Johnston, D. Delta signaling from the germ line controls the proliferation and differentiation of the somatic follicle cells during Drosophila oogenesis. Genes Dev 15, 1393–1405, doi:10.1101/gad.200901 (2001).

16. Gonzalez-Reyes, A. & St Johnston, D. Patterning of the follicle cell epithelium along the anterior-posterior axis during Drosophila oogenesis. Development 125, 2837–2846 (1998).

17. Keller Larkin, M., et al. Role of Notch pathway in terminal follicle cell differentiation during Drosophila oogenesis. Dev Genes Evol 209, 301–311, doi:10.1007/s004270050256 (1999).

18. Roth, S. Drosophila oogenesis: coordinating germ line and soma. Curr Biol 11, R779–781, doi:10.1016/s0960-9822(01)00469-9 (2001).

19. Poulton, J. S. & Deng, W. M. Cell-cell communication and axis specification in the Drosophila oocyte. Dev Biol 311, 1–10, doi:10.1016/j.ydbio.2007.08.030 (2007).

20. Xi, R., McGregor, J. R. & Harrison, D. A. A gradient of JAK pathway activity patterns the anterior-posterior axis of the follicular epithelium. Dev Cell 4, 167–177, doi:10.1016/s1534-5807(02)00412-4 (2003).

21. Kolahi, K. S. et al. Quantitative analysis of epithelial morphogenesis in Drosophila oogenesis: New insights based on morphometric analysis and mechanical modeling. Dev Biol 331, 129–139, doi:10.1016/j.ydbio.2009.04.028 (2009).

22. Balaji, R., Weichselberger, V. & Classen, A. K. Response of Drosophila epithelial cell and tissue shape to external forces in vivo. Development 146, doi:ARTN dev17125610.1242/dev.171256 (2019).

23. Alegot, H., Pouchin, P., Bardot, O. & Mirouse, V. Jak-Stat pathway induces Drosophila follicle elongation by a gradient of apical contractility. Elife 7, doi:10.7554/eLife.32943 (2018).

24. McInnes, L., Healy, J. & Melville, J. UMAP: Uniform Manifold Approximation and Projection for Dimension Reduction. *arXiv:1802.03426* (2018).

25. Timmons, A. K. et al. Phagocytosis genes nonautonomously promote developmental cell death in the Drosophila ovary. Proc Natl Acad Sci U S A 113, E1246–1255, doi:10.1073/pnas.1522830113 (2016).

26. Timmons, A. K., Mondragon, A. A., Meehan, T. L. & McCall, K. Control of non-apoptotic nurse cell death by engulfment genes in Drosophila. Fly (Austin) 11, 104–111, doi:10.1080/19336934.2016.1238993 (2017).

27. Chlasta, J. et al. Variations in basement membrane mechanics are linked to epithelial morphogenesis. Development 144, 4350–4362, doi:10.1242/dev.152652 (2017).

28. Grammont, M. Adherens junction remodeling by the Notch pathway in Drosophila melanogaster oogenesis. J Cell Biol 177, 139–150, doi:10.1083/jcb.200609079 (2007).

29. Lamire, L. A. et al. Gradient in cytoplasmic pressure in germline cells controls overlying epithelial cell morphogenesis. PLoS Biol 18, e3000940, doi:10.1371/journal.pbio.3000940 (2020).

30. Fletcher, G. C. et al. Mechanical strain regulates the Hippo pathway in Drosophila. Development 145, doi:10.1242/dev.159467 (2018).

31. Kaya-Copur, A. et al. The Hippo pathway controls myofibril assembly and muscle fiber growth by regulating sarcomeric gene expression. Elife 10, doi:10.7554/eLife.63726 (2021).

32. Hegde, R. S., Roychoudhury, K. & Pandey, R. N. The multi-functional eyes absent proteins. Crit Rev Biochem Mol Biol 55, 372–385, doi:10.1080/10409238.2020.1796922 (2020).

33. Jin, M. & Mardon, G. Distinct Biochemical Activities of Eyes absent During Drosophila Eye Development. Sci Rep 6, 23228, doi:10.1038/srep23228 (2016).

34. Rebay, I. Multiple Functions of the Eya Phosphotyrosine Phosphatase. Mol Cell Biol 36, 668–677, doi:10.1128/MCB.00976-15 (2015).

35. Bonini, N. M., Leiserson, W. M. & Benzer, S. The eyes absent gene: genetic control of cell survival and differentiation in the developing Drosophila eye. Cell 72, 379–395, doi:10.1016/0092-8674(93)90115-7 (1993).

36. Jevitt, A. et al. A single-cell atlas of adult Drosophila ovary identifies transcriptional programs and somatic cell lineage regulating oogenesis. PLoS Biol 18, e3000538, doi:10.1371/journal.pbio.3000538 (2020).

37. Rust, K. et al. A single-cell atlas and lineage analysis of the adult Drosophila ovary. Nat Commun 11, 5628, doi:10.1038/s41467-020-19361-0 (2020).

38. Chang, Y. C., Jang, A. C., Lin, C. H. & Montell, D. J. Castor is required for Hedgehog-dependent cell-fate specification and follicle stem cell maintenance in Drosophila oogenesis. Proc Natl Acad Sci U S A 110, E1734–1742, doi:10.1073/pnas.1300725110 (2013).

39. Fabrizio, J. J., Boyle, M. & DiNardo, S. A somatic role for eyes absent (eya) and sine oculis (so) in Drosophila spermatocyte development. Dev Biol 258, 117–128, doi:10.1016/s0012-1606(03)00127-1 (2003).

40. Demarco, R. S., Eikenes, A. H., Haglund, K. & Jones, D. L. Investigating spermatogenesis in Drosophila melanogaster. Methods 68, 218–227, doi:10.1016/j.ymeth.2014.04.020 (2014).

41. Moure, A. & Gomez, H. Phase-Field Modeling of Individual and Collective Cell Migration. Archives of Computational Methods in Engineering 28, 311–344, doi:10.1007/s11831-019-09377-1 (2021).

42. Nonomura, M. Study on multicellular systems using a phase field model. PLoS One 7, e33501, doi:10.1371/journal.pone.0033501 (2012).

43. Spradling, A. C. Developmental genetics of oogenesis. The development of Drosophila melanogaster, 1–70 (1993).

44. Jia, D., Xu, Q., Xie, Q., Mio, W. & Deng, W. M. Automatic stage identification of Drosophila egg chamber based on DAPI images. Sci Rep 6, 18850, doi:10.1038/srep18850 (2016).

